# Mapping viscosity in discrete subcellular locations with a BODIPY based fluorescent probe

**DOI:** 10.1101/522532

**Authors:** Lior Pytowski, Alex C. Foley, Zayra E. Hernández, Niall Moon, Timothy J. Donohoe, David J. Vaux

## Abstract

Numerous cellular processes, including enzyme behaviour, signalling, and protein folding and transport are highly influenced by the local microviscosity environment within living cells. Molecular rotors are fluorescent molecules that respond to the viscosity of their environment through changes in both the intensity and lifetime of their fluorescence. We have synthesised a novel benzyl-guanine derivatized boron-dipyrromethene (BODIPY) molecular rotor that is a substrate for the SNAP-tag targeting system (named BG-BODIPY), allowing us to target the rotor to discrete locations within the living cell. We demonstrate that BG-BODIPY reports viscosity, and that this can be measured either through fluorescence lifetime or intensity ratiometric measurements. The relative microviscosities within the ER, Golgi, mitochondrial matrix, peroxisomes, lysosomes, cytoplasm, and nucleoplasm were significantly different. Additionally, this approach permitted fluorescence lifetime imaging microscopy (FLIM) to determine the absolute viscosity within both mitochondria and stress granules, showcasing BG-BODIPY’s usefulness in studying both membrane-bound and membraneless organelles. These results highlight targeted BG-BODIPY’s broad usefulness for making measurements of cellular viscosity both with FLIM and conventional ratiometric confocal microscopy, the latter option greatly extending the accessibility of the technique although limited to relative meassurements.

**Statement of Significance:** Local viscosity affects molecular behaviour from diffusion and conformational changes to enzyme kinetics and has important implications for cell and tissue function. Mechanical methods for measurement of viscosity average over large volumes and long times and are thus unsuitable for rapid changes on small scales that are biologically relevant. This paper reports a novel optical fluorescence method using genome edited cells to deliver a viscosity reporter to tightly defined locations inside living cells, from which non-destructive organelle-specific measurements can be repeatedly made. The local viscosity of seven separate organelles in living cultured human cells is shown for the first time, together with the viscosity behaviour of a membraneless organelle as it is induced in cells by stress.

## Introduction

Viscosity is an essential biophysical factor influencing numerous cellular processes for which diffusion has evolved to become the rate-limiting factor, such as signal transduction, protein-protein interactions, and intracellular transport of macromolecular cargo^1^. Traditional methods of determining the viscosity of a substance by measuring its resistance to shear or tensile stress operate on continuous material at far too large scale to appreciate the microviscous environment within a cell. As we begin to understand that single cellular compartments may have significant local variations within them, and that our view of cellular compartmentalisation is lent further complexity by the presence of membraneless organelles, it becomes increasingly clear that we require a means of measuring viscosity at discrete locations in living cells^2^.

Most attempts at measuring microviscosity in living cells have relied primarily upon a class of synthetic fluorescent molecules that respond to the local viscous environment, termed molecular rotors^3^. The viscosity-dependence of these fluorophores is mediated by the presence of a hinge domain in the probes that allows intramolecular rotation, creating competition for releasing the energy of an absorbed photon through either fluorescence or non-radiative decay via mechanical twisting of the molecule. Among this class of fluorophores, the most commonly utilised are those which can access Twisted Intramolecular Charge Transfer (TICT) states, which are states of lower energy charge transfer resulting in decreased fluorescence emission intensity^1^. The fluorescence intensity is directly influenced by the local viscosity, as an increase in viscosity will reduce the amount of intramolecular twisting, forcing the energy emission to occur through fluorescence, and vice versa. This results in both an increase in the fluorescence intensity and fluorescence lifetime of the molecule as the local microviscosity increases.

Boron-dipyrromethene (BODIPY) is one of the most commonly used molecular rotors. BODIPY belongs to the TICT class of molecular rotors, and is therefore a viscosity reporter in that its de-excitation after absorbing a photon occurs through a competition between fluorescence and non-radiative decay through intramolecular rotation^3,4^. BODIPY molecular rotors have a number of attractive traits, such as their strong UV absorbance, their stability in physiological conditions, and their tunability with small chemical modifications^5,6^.

There has been a spate of efforts to map eukaryotic cells in terms of viscosity. Kuimova et al. (2008)^4^ loaded SK-OV-3c cells with *meso*-substituted 4,4′-difluoro-4-bora-3a,4a-diaza-*s*-indacene not conjugated to any targeting molecule, measuring the viscosity in various subcellular localisations. Levitt et al. (2009)^6^ used two meso-substituted BODIPY based fluors conjugated to hydrophobic groups designed to drive indiscriminate incorporation into the hydrophobic interior of non-aqueous cellular membranes, and discovered that the intracellular environment contains a wide range of viscosities. Several groups have targeted boron-dipyrrin (BODIPY) modified by the addition of triphenylphosphonium to target the dye to the mitochondria with varying degrees of success^7,8^. More recently, Chambers et al. (2018)^9^, have developed a chloroalkane-BODIPY capable of targeting to specific organelles via a transiently expressed HALO-tag protein targeted to mitochondria or the ER lumen.

In parallel, our group has developed a method for targeting BODIPY conjugated via a linker moiety to a benzylguanine group to discrete cellular locations through SNAP-tag technology^10^. SNAP-tag utilises an *O*^6^-alkylguanine-DNA alkyltransferase enzyme expressed either through transient expression or CRISPR/Cas-9 knock-in. This modified enzyme binds a benzylguanine group to produce a covalent adduct that cannot be resolved, so that the enzyme becomes covalently linked to this substrate. Our method additionally benefits from the expression of an mCherry fluorophore, the SNAP-tag enzyme and the localisation signals, allowing us to map numerous cellular locations with a single BODIPY-based probe through both fluorescence lifetime imaging microscopy (FLIM) and ratiometric imaging.

In this paper we demonstrate that a novel SNAP-tag compatible BODIPY probe can be targeted to both membrane-bound subcellular compartments and membraneless organelles. We show that this new probe reports microviscosity in discrete cellular locations through both FLIM and ratiometric imaging when the BODIPY fluorophore is bound to a targeted SNAP-tag-mCherry fusion protein. This system presents many possibilities, as it can be combined with any number of target sequences or proteins, and can accommodate numerous sensor fluorophores (for example for calcium ion concentration, pH, and redox potential)^11–13^ so long as they can be conjugated to a benzylguanine group and exhibit cell-penetrant properties. Further, the ability to measure viscosity by quantitating either the intensity or the lifetime of the fluorophore signal confers upon this technique an improved degree of flexibility, with each complementary method exhibiting advantages over the other.

## Materials and methods

### BG-BODIPY synthesis

BG-BODIPY synthesis is fully described in supplementary materials.

### Fluorescence spectra determination

Spectra were measured in water on a Fluorolog FL3-22 spectrofluorimeter using FluorEssence software v3.5 (HORIBA Jobin Yvon), with a xenon lamp, 0.1 s dwell time and 10 repeats at room temperature. Scan slit was 3.00 nm and a TCC detector was used.

### Cell culture

HeLa cells (from lab stock, traceable to ATCC) were cultured in Dulbecco’s Modified Eagle’s Medium (DMEM) with 10% foetal bovine serum (FBS, Sigma F4135), 1% non-essential amino acids, and penicillin and streptomycin (100 units/ml). Cells were incubated at 37°C with 5% CO_2_ in a humid atmosphere. For FLIM and confocal imaging cells were plated the day before, or three days prior to in the case of transfection, imaging on optical bottomed 96-well plates (PerkinElmer).

### Drug treatment and staining of cultured cells

#### MitoTracker staining

Cells were stained with MitoTracker Deep Red 633 (Invitrogen M22426) following manufacturer’s recommendations.

#### Zinc chloride treatment

TDP-43 aggregation was induced with complete DMEM supplemented 100 µM ZnCl_2_ (Sigma, Z0152) overnight. ZnCl_2_ stock was prepared in water.

### Generation of cell lines by CRISPR/Cas9

#### Donor plasmid generation

mCherry-SNAP-tag with the appropriate localisation signals and bGH poly(A) signal was inserted in the *AAVS1* locus (GenBank accession AC010327.8 (7774 - 11429)) using a double-nicking strategy with the Cas9 D10A nickase mutant^14^. The donor plasmid consisted of the insert enclosed in an 840 bp C-term homology arm GenBank accession AC010327.8 (8372 - 9211) and an 809 bp N-term homology arm GenBank accession AC010327.8 (9212 - 10021) between position 70 and 71 of the lacZa gene of pBluescript. The donor plasmids were assembled by isothermal assembly (NEBuilder® HiFi DNA Assembly).

#### Localisations signals

Localisation signals were based on the protein localization signals database LocSigDB^15^ and Chertkova et al. 2017 findings^16^.

#### gRNA plasmid generation

gRNA and plasmid design were done following Ran et al. (2013)^17^ recommendations. Guides were designed with MIT design tool: http://crispr.mit.edu/ and inserted into Addgene Plasmid #48141 pSpCas9n(BB)-2A-Puro (a gift from Feng Zhang, MIT, USA). Guides RNA were 5’GTCCCTAGTGGCCCCACTGT3’ and 5’GACAGAAAAGCCCCATCCTT3’.

#### Transfection and cell sorting

Plasmids containing the guides and donor templates were transfected in HeLa cells using Lipofectamine 2000 (Invitrogen, 11668030) following manufacturer recommendations. Cells were then left to recover and to express the fusion protein for a week before proceeding with cell sorting. Cell sorting was done on a BD FACS Aria III run with the Diva software version 8.0.2. Positive cells were sorted if red fluorescence was 10^3^ times brighter that control cells (excitation at 561nm 50mW, emission filter 610/20). Positive cells were cloned by single cell sorting into an optical 96-well plate containing preconditioned media and the expected localisation of the fusion protein was confirmed by confocal microscopy.

### Plasmids

Lysosome localisation signal (residues 27 to 407 of Human LAMP1) was extracted by PCR from Addgene’s plasmid #55308. Golgi localisation signal (residues 3131 to 3259 of Human Giantin) was extracted by PCR from Addgene’s plasmid #85048. The mitochondrial localisation signal (Residues 1 to 29 of Human COX8, 4 times)^18^ was extracted by PCR from Addgenes’s plasmid #98819. The SNAP-tag coding sequence was extracted from plasmid #N9183S New England Biolabs (Hitchin, Herts, UK).

The control plasmid pSNAPf-Cox8A for transient targeted SNAP-tag enzyme expression in mitochondria was obtained from New England Biolabs (#N9185). TDP43 was inserted by Gibson assembly at the C terminus of SNAP-tag (tdp43-EGFP was a gift from Zuoshang Xu, Addgene plasmid # 28198). Plasmids were delivered to cells using Lipofectamine 2000 according to the manufacturer’s instructions. Cells were labelled with BG-BODIPY and imaged as described below after allowing 24 hours post-transfection for protein expression.

### Live cell time course

Cells were plated in 24 well plates then post adhesion treated with 2.5 µM BG-BODIPY for 60 minutes. Cells were then washed three times for 10 minutes with 5% delipidated BSA in DMEM in order to back extract the remaining free fluor. Following washing, cells were covered in culture medium for imaging. Cells were then imaged in an IncuCyte ZOOM (Sartorius) placed in a tissue culture incubator (37°C with 5% CO_2_ in a humid atmosphere) for 18h every 5 minutes using a 20x objective.

### SNAP-tag labelling for FLIM and confocal microscopy

Plated cells were treated with 2.5 µM BG-BODIPY for 60 minutes. Cells were then washed three times for 10 minutes with 5% delipidated BSA in DMEM. Following washing, cells were covered in Fluorobrite DMEM (ThermoFisher) medium with 10% FBS, Penicillin-Streptomycin for imaging.

### Fluorescence lifetime measurements

FLIM was carried out on an Olympus FV 1000 equipped with a Becker & Hickl SPC-150 Time-Correlated Single Photon Counting (TCSPC) module. A solid state 488 nm laser pulsed at 80 MHz was used as an excitation source. Fluorescence decay measurements were collected using the SPCM software and analysed using the SPCImage software provided by Becker & Hickl. A two exponential decay model was used for fitting the curves, denoted by the general equation (1):

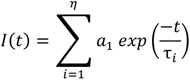

where *I(t)* is the time-independent fluorescence intensity, **τ** is the lifetime, α is the amplitude, and η is the number of decay coefficients.

All images were acquired with a Plan-Apochromat 60x objective to negate any polarization effects, as NA = 1.4 oil-immersion microscope lenses are free of anisotropy^19^. All measurements were performed at room temperature.

Following multi-exponential fitting, the raw lifetime data for each pixel was exported to Fiji^20^, where regions of interest were selected to create region specific lifetime histograms that were converted to cP using the equation log(Y)=(0.3035)X+2.581 where Y is the lifetime (ps) and X is the viscosity (cP).

### Confocal microscopy

Samples were imaged on a Zeiss LSM880 with Airyscan using a Plan-Apochromat 40x 1.3 NA oil immersion lens. For ratiometric imaging 16-bit images were acquired with Zen Black software. Channel registration was done using 0.1 µm TetraSpeck™ Microspheres (Invitrogen T7279) and images were corrected for translation, rotation and scaling in Zen Blue. Overlap coefficient was calculated with JACOP^21^. Ratiometric calculation was done by dividing the BODIPY image by the mCherry image (16-bit input images creating a 32-bit output) and the histogram was created with the “Histogram” tool (512 bins, ranging from 0.5 to 3) and the data was exported. Data was plotted in R with ggplot2 or in GraphPad Prism 8. Mitochondria line profiles were created by drawing a linear region of interest (ROI) perpendicular to the mitochondria long axis and the intensity across those ROIs were plotted with the “Plot Profile” tool. The line scans were then aligned to generate the averages and plotted with GraphPad Prism 8.

### Viscosity probe calibration

The Förster-Hoffmann equation (2) characterizes the power-law relationship of quantum yield and viscosity:

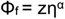

where *z* and α are constants and Φ_f_ is the fluorescence quantum yield. The fluorescent quantum yield, and the other main parameter of fluorescence, the fluorescence lifetime, are defined by (3)

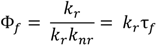

where *k_r_* and *k_nr_* are the radiative and non-radiative decay constants, respectively. It follows that a plot of log Φ_f_ against log η yields a straight line with slope α^22^.

To determine the effects of pH on the BODIPY probes, we created a series of MOPS buffers with pH’s that varied as integers from 4 to 10. These buffers were then each mixed with a series of glycerol volumes to create buffers ranging in viscosity ranging from 0% to 95% glycerol. These buffers with varying pH and glycerol concentration were pipetted into microfluidic chamber grids, as described below, by an iotaSciences microfluidics printer, along with unbound BG-BODIPY at a concentration of 2.5 µM. The lifetime and intensity of BG-BODIPY in these chambers were then measured by FLIM.

### Creation of the microfluidic chamber grids

An iotaSciences microfluidics printer^23^ was used to create 7 x 15 grids for the calibration of the BODIPY dyes. The bottom of a 60 mm tissue culture polystyrene dish (Corning) was covered with DMEM + 10% FBS and the excess media removed. This was overlaid with an immiscible liquid, FC-40 (Fluorinert™). At the microscale, gravitational and buoyant effects become negligible and the interfacial forces pin the aqueous phase under the FC-40 to the plastic of the dish. The hydrophobic stylus with a conical tip made of polytetrafluoroethylene (Teflon) of the iotaSciences microfluidics printer was then lowered through both liquids until it makes contact with the dish. FC-40 wets the Teflon tip better than water, and so as the stylus was dragged laterally, the DMEM was displaced by the FC-40, which becomes pinned to the surface of the dish. A grid with 1.9 mm x 1.9 mm chambers was drawn in such a fashion and the DMEM was then removed from each grid chamber, pipetted out by the iotaSciences printer, and replaced with a mixture containing 2.5 µM BODIPY and varying concentrations of glycerol (0 to 95%) and MOPS buffers with pHs ranging from 4 to 10.

### Electron Microscopy and analysis

Cells were trypsinised and high pressure frozen as for Johnson *et al* (2015)^24^, using a Leica EM ICE HPF. Samples were transferred to a Leica AFS2 freeze substitution unit and freeze substituted at −130°C in 0.2% uranyl acetate and 5% water in acetone, then infiltrated with HM20 Lowicryl resin as for Johnson *et al* (2015) and polymerised with UV light for a total of 48 h with the following schedule: −45°C for 24 h, warming to 0°C over 12 h, then for a further 12 h at 0°C. Ultrathin (90 nm) resin sections were taken from polymerised blocks using a Diatome diamond knife on a Leica Ultracut 7 and transferred to formvar coated 50 mesh nickel grids.

On-section immunolabelling was performed by serial transfer of grids between 50 ul droplets of solution at room temperature: Grids were placed, section side down, on a droplet of blocking buffer (1% BSA, 0.5% FCS, 1% Fish Gelatin in PBS) for 15 min and then incubated in rabbit anti-mCherry primary antibody (Abcam, ab167453) diluted 1:50 in blocking buffer for 30 min. Grids were washed in blocking buffer 5x 2 mins and then incubated in 10 nm colloidal gold-conjugated anti-rabbit secondary antibody (Abcam, ab27234) diluted 1:20 in blocking buffer for 30 min. Grids were washed as before, then washed 3x 2min in PBS and 3x 1min in Milli-Q water before being post-stained for 10 min with 2% UA, washed 5x 1 min with warm degassed water and then 10 min with Reynold’s lead citrate and washed 5×1 min with warm degassed water, then air dried.

Grids were imaged on a FEI Tecnai 12 TEM operated at 120 kV using a Gatan OneView camera. Images containing gold particles were segmented using Ilastik 1.3.3 following pixel classification and object classification protocols^25^. For pixel classification all colour/intensity, edge and texture features were selected. A parallel random forest classifier was used (VIGRA). Probability predictions were created and used as input for object classification. Pixels with a probability score above 0.6 were considered as contributing to gold particles if the whole object segmented was between 60 and 160 pixels (Supplementary figure 3).

## Results

### BG-BODIPY, a novel molecular rotor, reports microviscosity in live cells in both the lifetime and intensity domains

To measure microviscosity in discrete subcellular compartments, we created a viscosity sensor able to be targeted to multiple locations through a general mechanism. A BODIPY probe was synthesised to be used in conjunction with SNAP-tag technology, which we termed benzylguanine-BODIPY, or BG-BODIPY (Figure 1A). The synthetic approach is fully described in Supplementary material. The maximal absorbance of BG-BODIPY is at 498 nm and the emission peak is at 515 nm (Figure 1B). The probe comprised a BODIPY molecular rotor conjugated via a linker to a benzylguanine group, a substrate for the modified O^6^-alkyguanine-DNA alkyltransferase (AGT) employed by SNAP-tag, altered such that upon attempting to cleave its substrate, the two become covalently linked^10^. We then targeted this BG-BODIPY probe in living cells with a high degree of specificity by expressing modified SNAP-tag as a fusion protein targeted to specific organelles.

**Figure 1:**
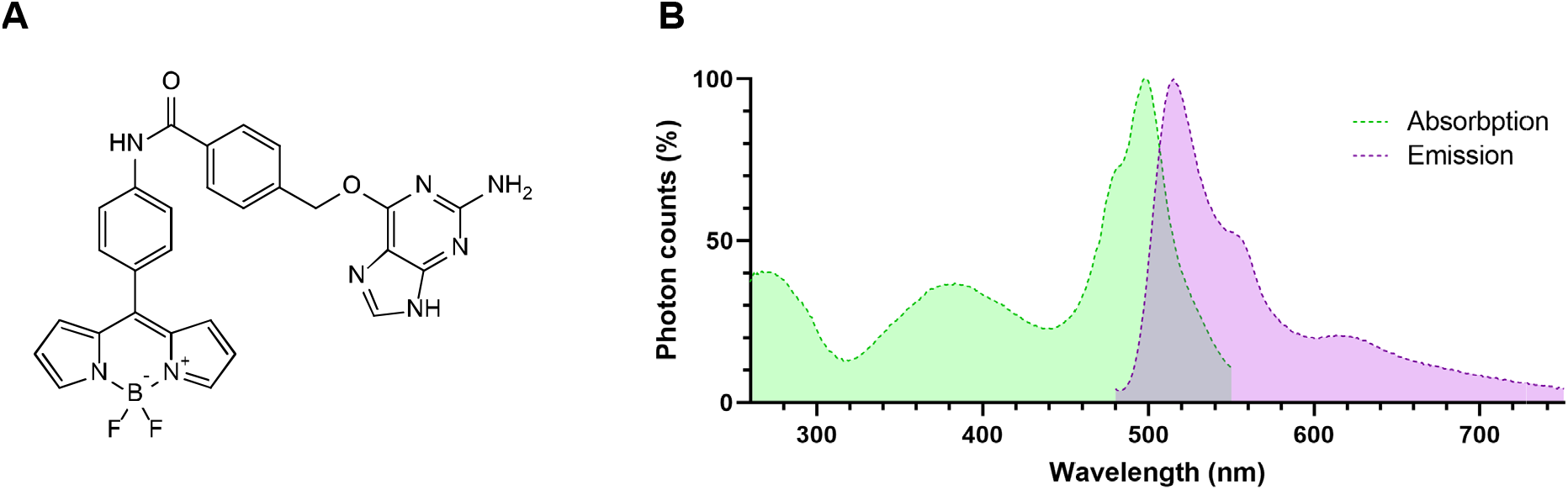
BG-BODIPY structure and spectra. **A.** Structure of BG-BODIPY. BODIPY molecular rotor conjugated via a linker to a benzylguanine group. **B.** Absorption and emission spectra of BG-BODIPY.

In order to ensure that BG-BODIPY still responded to viscosity and that this response occurred over a useful and measurable range of lifetimes, we used an iotaSciences microfluidics printer^23^ to create grids of 2 mm fluid cells, partitioned with a water-immiscible fluid (FC-40), and containing varying percentages (vol/vol) of glycerol (0 to 95%) and MOPS buffer at 5 different integer pH values (4 to 8). Supplementary figure 1 shows lifetime images from this experiment together with a demonstration of the software analysis that confirms a single lifetime for the images. We found that the pH did not have a significant effect on either the measured lifetime or intensity of BG-BODIPY across the whole range of viscosities (Figure 2A, C). At each pH, we plotted the log Φ_f_ against log η and determined that the ratio followed a linear relationship (Figure 2B). Furthermore the lifetime and intensity are proportional over a range of pH and viscosity values (Figure 2D).

**Figure 2:**
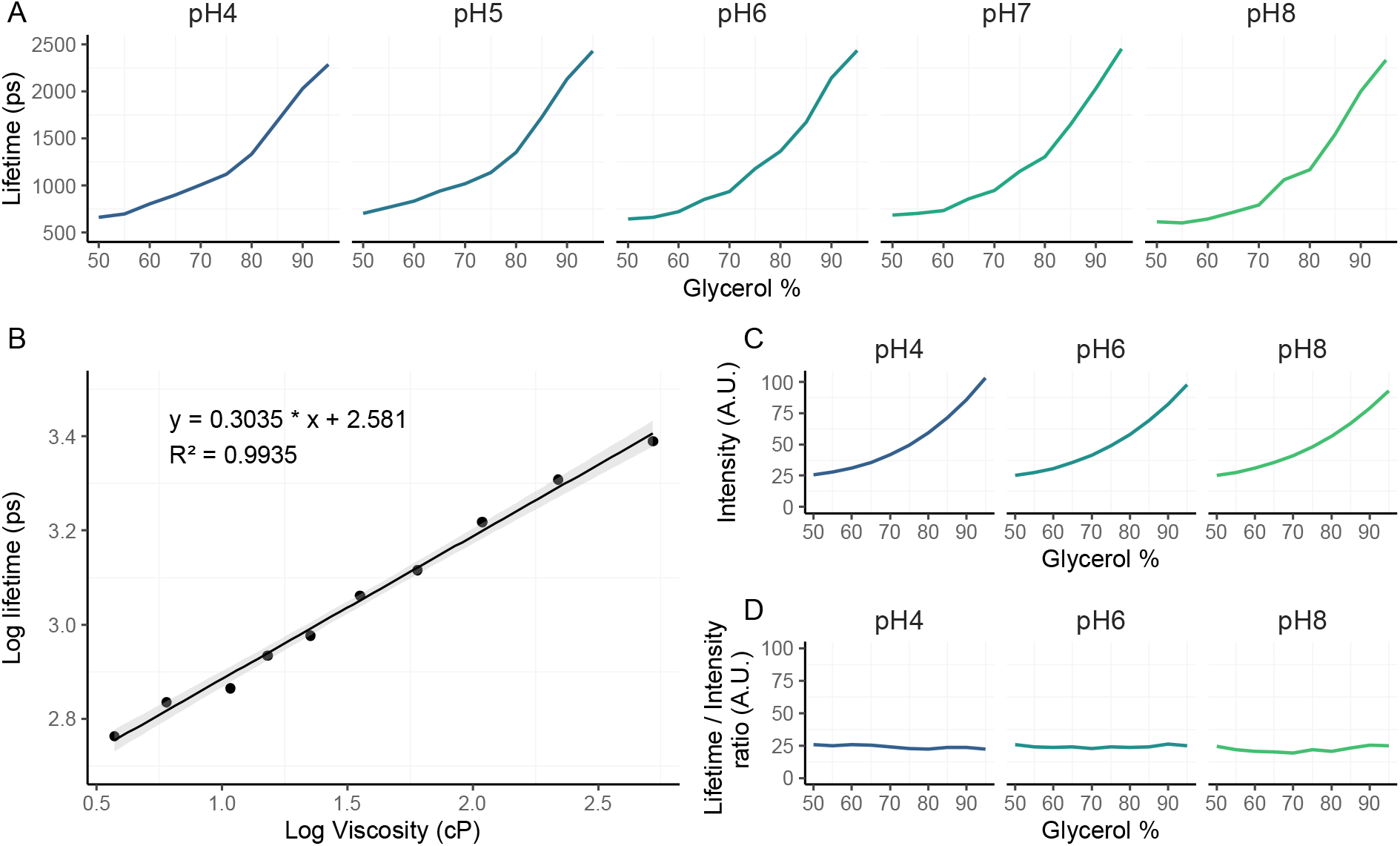
BG-BODIPY reports microviscosity in both the lifetime and intensity domains and is pH insensitive. Calibration of BG-BODIPY. **A**. The lifetime of BG-BODIPY in picoseconds (Y axis) measured at various pH values and glycerol concentrations (X axis). Curve fitting with a sigmoid model yielded Hill coefficients that did not differ significantly between different pHs (p=0.70). **B**. The log plot of fluorescence lifetime as a function of the log viscosity for BG-BODIPY in pH 7 MOPS buffer. The function is linear, expressed by the equation Y=(0.3035)X+2.581. **C**. The intensity of BG-BODIPY fluorescence measured at various pH values (y axis) and glycerol concentrations (x axis). Curve fitting with a sigmoid model yielded Hill coefficients that did not differ significantly between different pHs (p=0.61). **D**. The ratio of lifetime over intensity at various pH values (y axis) and glycerol concentrations (x axis). Curve fitting with a sigmoid model yielded Hill coefficients and Y intercepts that did not differ significantly between different pHs (p=0.23 and p=0.27 respectively). Supplementary figure 1 contains sample images used for the calibration curves and a sample probability histogram derived from the SPCImage software.

### BG-BODIPY can be targeted to discrete membrane-bound organelles

In order to target the BG-BODIPY to specific organelles, we expressed SNAP-tag fusion proteins either through transient transfection or using CRISPR-Cas9 to knock-in the sequence into the *AAVS1* safe harbour genomic locus in human cells. Initial experiments revealed significant non-specific dye retention after cell exposure, but this background could be eliminated by back-extraction using three washes for 10 minutes with 5% delipidated bovine serum albumin (BSA) in DMEM as an absorber for the excess probe (supplemental figure 2).

Once we had a means of back-extracting the untargeted BG-BODIPY, we compared the co-localisation of Mitotracker with BG-BODIPY stained cells expressing a transiently transfected Cytochrome c oxidase 8A (Cox8) SNAP-tag fusion protein, or stably expressing a SNAP-tag fusion with mCherry and a mitochondrial targeting sequence after CRISPR/Cas-9 knock in of the appropriate construct to the *AAVS1* safe harbour locus. We observed an almost complete overlap of the BG-BODIPY and Mitotracker signals for both the CRISPR cell lines and the transiently transfected cells (Figure 3 A and B).

**Figure 3:**
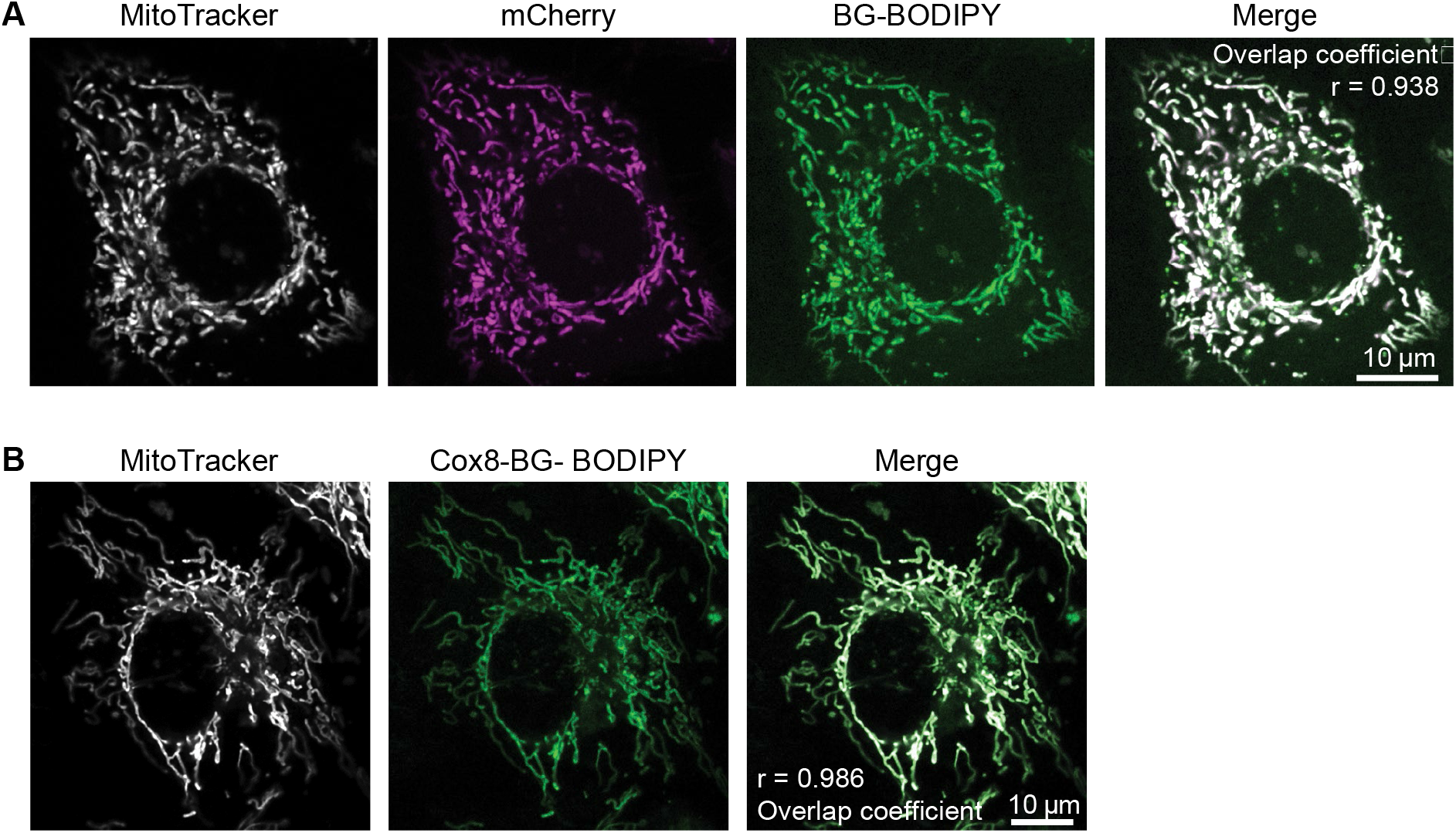
BG-BODIPY targets specifically to subcellular compartments. Colocalization of Mitotracker with **A**. mCherry and BG-BODIPY targeted to the mitochondria via residues 1 to 29 of Human COX8 (Overlap Coefficient=0.938 r²=0.879) and **B**. transient expression of a Cox8-SNAP-tag fusion protein (Overlap Coefficient=0.986 r²= 0.971).

For ultrastructural confirmation of the targeted delivery of the fusion proteins to the desired organelles we then performed colloidal gold immunolabeling for mCherry on ultrathin sections from the CRISPR/Cas-9 edited cells, followed by transmission electron microscopy (Figure 4). This approach confirmed that the mCherry moiety of the fusion protein was exposed within the interior of the targeted compartments (Figure 4).

**Figure 4:**
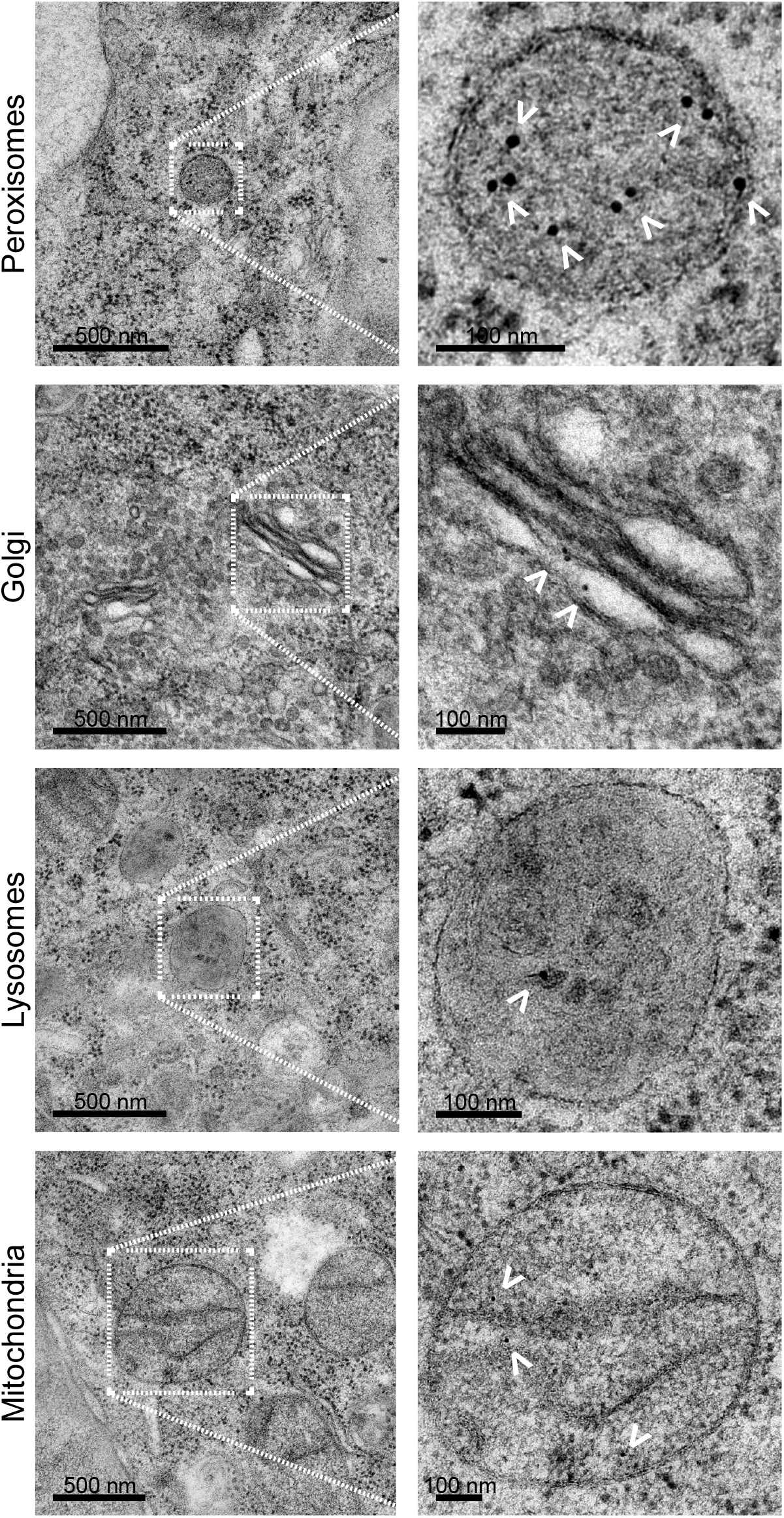
mCherry-BG-BODIPY can be targeted to all cellular compartments. The mCherry-SNAP-tag fusion protein targeted to distinct cellular compartments was constitutively expressed from the *AAVS1* locus. Cells were fixed and prepared for transmission electron microscopy by immuno-gold staining with anti-mCherry antibody. White arrowheads indicate colloidal gold particles. Note that these can be distinguished from other particles such as glycogen granules in the cytoplasm by their extreme electron density.

### BG-BODIPY does not affect cell viability

Transient expression of high levels of fusion proteins from transfected plasmids bearing high activity viral promoters can saturate normal biosynthetic pathways and lead to aberrant localisation of the fusion protein, progressive aggregate accumulation and severe cellular toxicity. Having established that the lower expression levels yielded by insertion into the *AAVS1* safe haven locus led to appropriately targeted fusion protein, with no saturation or aberrant localisation, we then turned our attention to cytotoxicity. In order to demonstrate that neither the expression of the fusion protein nor the subsequent loading of the SNAP-tag moiety with BG-BODIPY caused cellular cytotoxicity we collected time-lapse images at five-minute intervals for eighteen hours after labelling of each of the cell lines. We found that BG--BODIPY exposure and the back-extraction do not affect cell viability as cells continue to divide and behave normally (Movie 1).

**Movie 1: BG-BODIPY does not affect cell viability**. Cells were treated with BG-BODIPY as described in the methods and were imaged in an IncuCyte for 18h every 5 minutes. From left to right, top to bottom: WT, ER, Golgi apparatus, Lysosomes, Peroxisomes, Mitochondria, Nucleoplasm, Cytoplasm mCherry-SNAP-tag tagged cells lines.

### mCherry-BG-BODIPY can be targeted to selected cellular compartments and reports relative microviscosity

To demonstrate the usefulness of BG-BODIPY in mapping viscosity throughout the cell, we measured the ratio of BG-BODIPY to mCherry fluorescence in all of the knock-in lines, which targeted the SNAP-tag enzyme to the endoplasmic reticulum (ER), the Golgi apparatus, lysosomes, peroxisomes, mitochondria, the nucleoplasm, and the cytoplasm (Figure 5). We observed viscosities that vary significantly between the various membrane-bound organelles. For instance, the mean ratio value increased from the ER (1.67) to the Golgi (1.76) (p<0.001), but the range of the ratio values broadened considerably. The second highest mean ratio, after the Golgi, was in the mitochondrial matrix (1.70). The lowest of the viscosities reported was in the nucleoplasm (1.45) which was significantly lower than the cytoplasm (1.56) (p<0.001). Lysosomes had a mean ratio slightly less than that of the cytoplasm (1.52), while peroxisomes had an intermediate mean ratio (1.62) but exhibited by far the largest variance.

**Figure 5:**
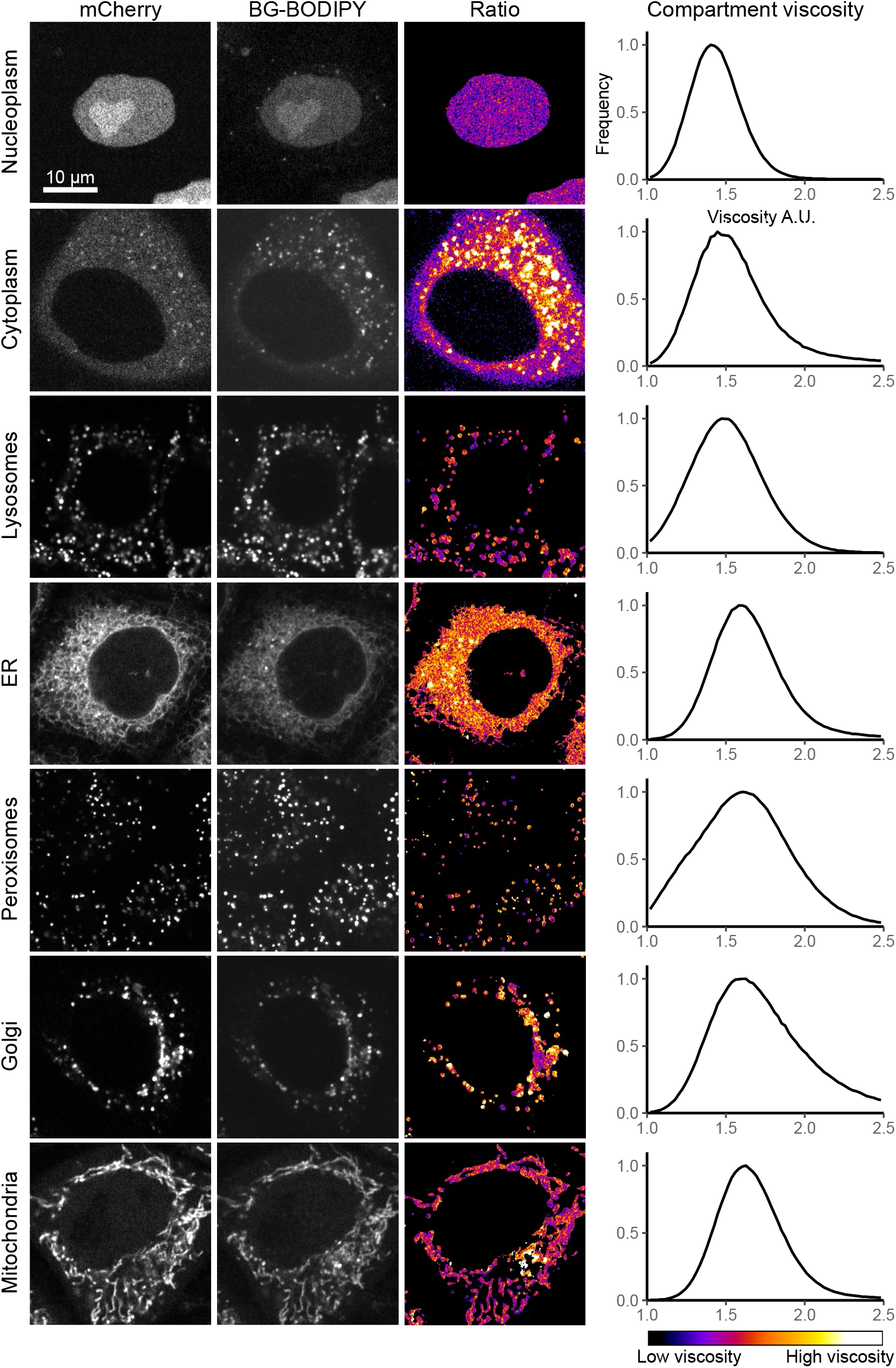
mCherry-BG-BODIPY can be targeted to all cellular compartments and reports relative microviscosity. The viscosity is highly variable in different cellular compartments. The mCherry-SNAP-tag fusion protein targeted to distinct cellular compartments was constitutively expressed from the *AAVS1* locus and tagged with BG-BODIPY by subsequent incubation with the probe. Ratiometric analysis of the fluorescence intensity of mCherry and BODIPY enabled us to assess the compartment viscosity. The analysis shown is based on 3 experimental replicates of approximately 40 to 50 cells each.

### Ratiometric measurements are composites of two-component Gaussian mixtures

We hypothesised that the frequency distribution of pixel ratios is composed of two populations, one of which is the BODIPY bound to SNAP-tag and another that is the BODIPY that aggregated within the cell. To assess this hypothesis, the frequency distribution of the ratios was fitted to a single or double component Gaussian mixture model. We found that although all distributions are better fitted by two gaussian mixtures, the ratios for the peroxisomes, lysosomes and nucleoplasm have only a minimal reduction in the sum of squared differences (SSD) between the predicted and actual y values (0.007 to 0.022 of difference) while the remaining compartments had a 0.064 to 0.414 SSD improvement when adding a second component (Figure 6). This analysis may help to identify regions of high ratio that may correspond to aggregates visible in the image.

**Figure 6:**
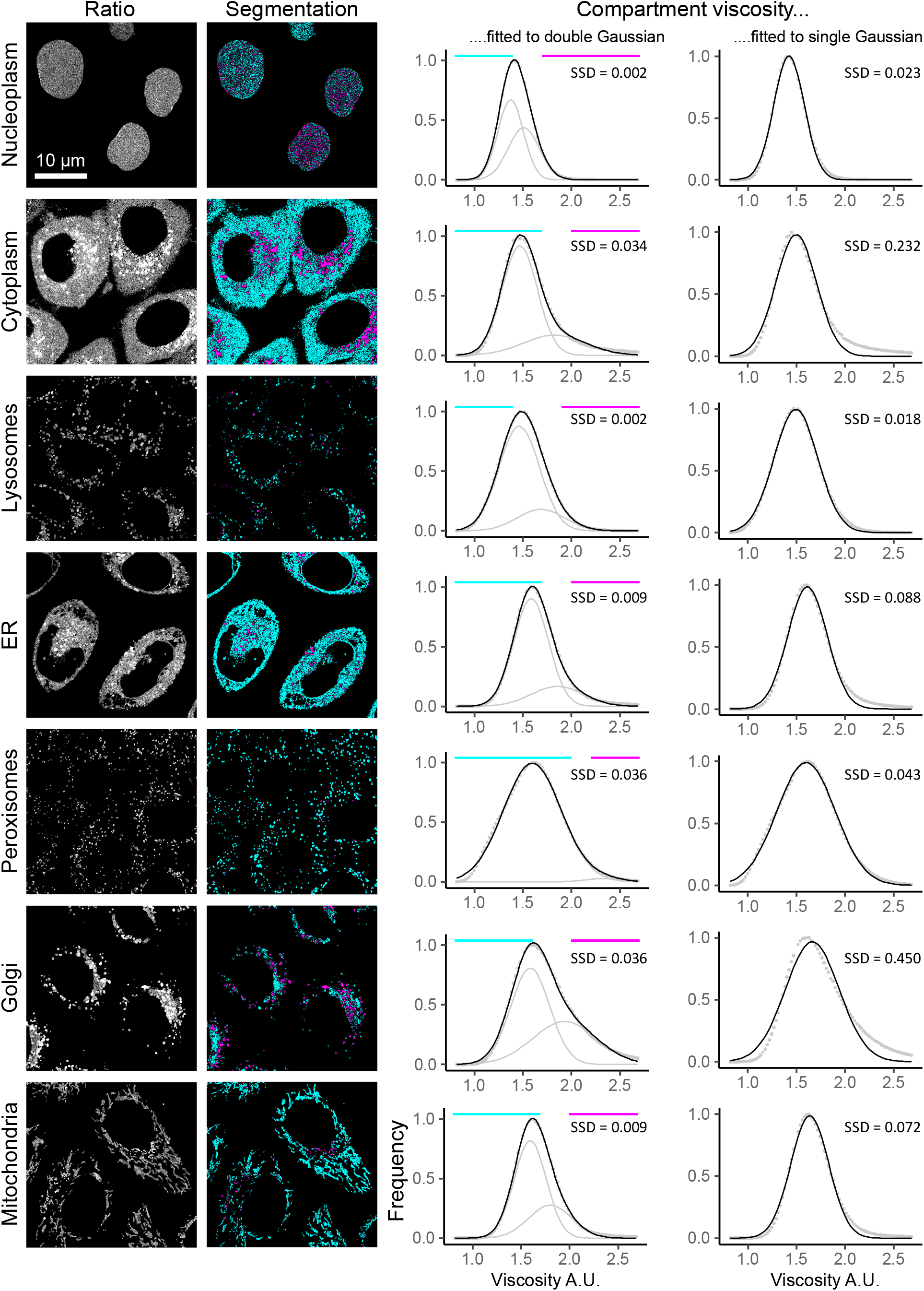
Ratiometric measurements are composites of two-component Gaussian mixtures. Single or double gaussian mixtures were fitted to the ratiometric distributions. Cyan and magenta regions in righthand segmented images correspond to predominantly the first lower viscosity (left) or second higher viscosity (right) gaussian component. SSD: Sum of Squared Differences between model (solid black line) and observed data (dotted line).

### Ratiometric measurements for some compartments can be approximated to cP values

To assess whether ratiometric measurements could be equated to cP values we measured the viscosity in the targeted compartments by FLIM. Equipped with the cP values plus the ratiometric measurements for each compartment we correlated them and found a simple monotonic correlation between ratio and cP in the cytoplasm, Nucleus, Lysosomes, Peroxisomes, Mitochondria and ER. Only the Golgi apparatus value lies outside the 95% confidence interval. We do not know the reason for this but does not seem to be due to increased heterogeneity of viscosity within the organelle (because variance is not increased). It cannot be accounted for by differential maturation of the mCherry and SNAP-tag moieties since the Golgi targeted probe has been delivered via the ER (values for which lie on the calibration curve). It is unlikely to be due to a mCherry brightness reduction caused by low pH because if this were the case the lysosomes would also suffer from it. Note that the Golgi apparatus is site of segregation of contents both during cargo processing, and selection/concentration of vesicular content; this could give rise to regions of differing viscosity.

### BG-BODIPY reports microviscosity in organelle sub-compartments

In order to test subcompartment-specific reporting of viscosity, we turned to the two mitochondrial targeted SNAP-tag probes. In the CRISPR/Cas9 engineered cells the mCherry-SNAP-tag fusion protein is targeted to the mitochondria via residues 1 to 29 of Human COX8, and is delivered to the mitochondrial matrix^18^ (Table 1). On the other hand, the transiently expressed mitochondrial targeting vector pSNAPf-Cox8A encodes the entire Cytochrome C oxidase, subunit (COX8A, UniProtKB - P10176) as a protein fusion to the N-terminus of the SNAP-tag. Cytochrome C oxidase is located in the inner mitochondrial membrane, with its C-terminus in the intermembrane space, such that the SNAP-tag enzyme is then delivered to the intermembrane space^26^ (Figure 8A). Confirmation of these two different sub compartment localisations is shown by averaged fluorescence intensity in line-scans taken orthogonal to the long axis of labelled mitochondria in living cells after BG-BODIPY labelling (Figure 8B). Label in the matrix space shows a single peak, while label in the intermembrane space shows the expected two peaks, although both probes show almost complete overlap with a Mitotracker signal (Figure 3). FLIM measurements reveal that the local environment of the BG-BODIPY in these two locations is strikingly different (Figure 8C), with a much higher viscosity in the matrix space than in the intermembrane space.

**Figure 7:**
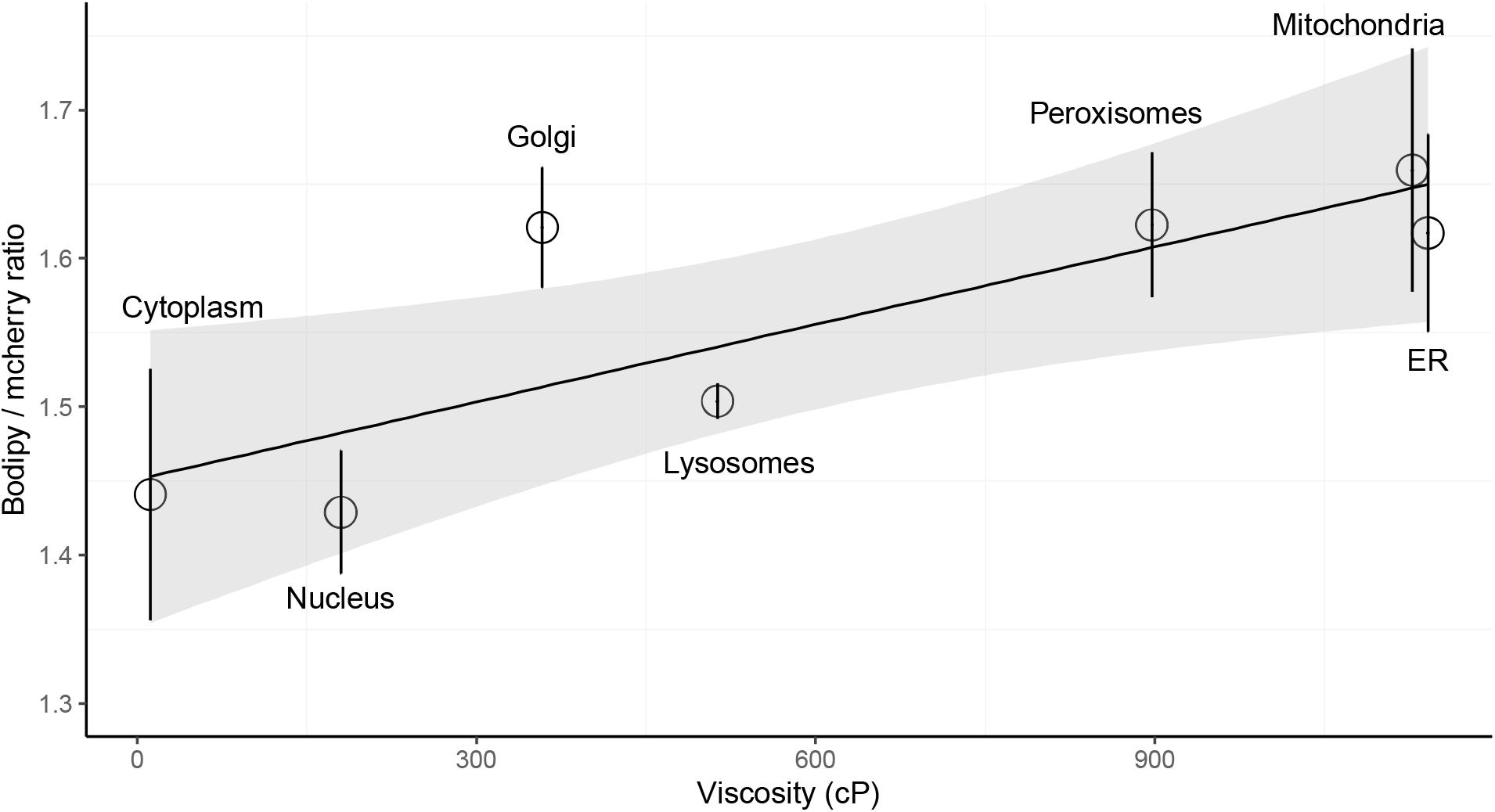
Ratiometric measurements for some compartments can be approximated to cP values. The average ratio (Y axis) for each compartment is plotted against the measured cP viscosity by FLIM (X axis). Vertical lines represent SEM. Grey shaded area represents 95% confidence intervals.

**Figure 8:**
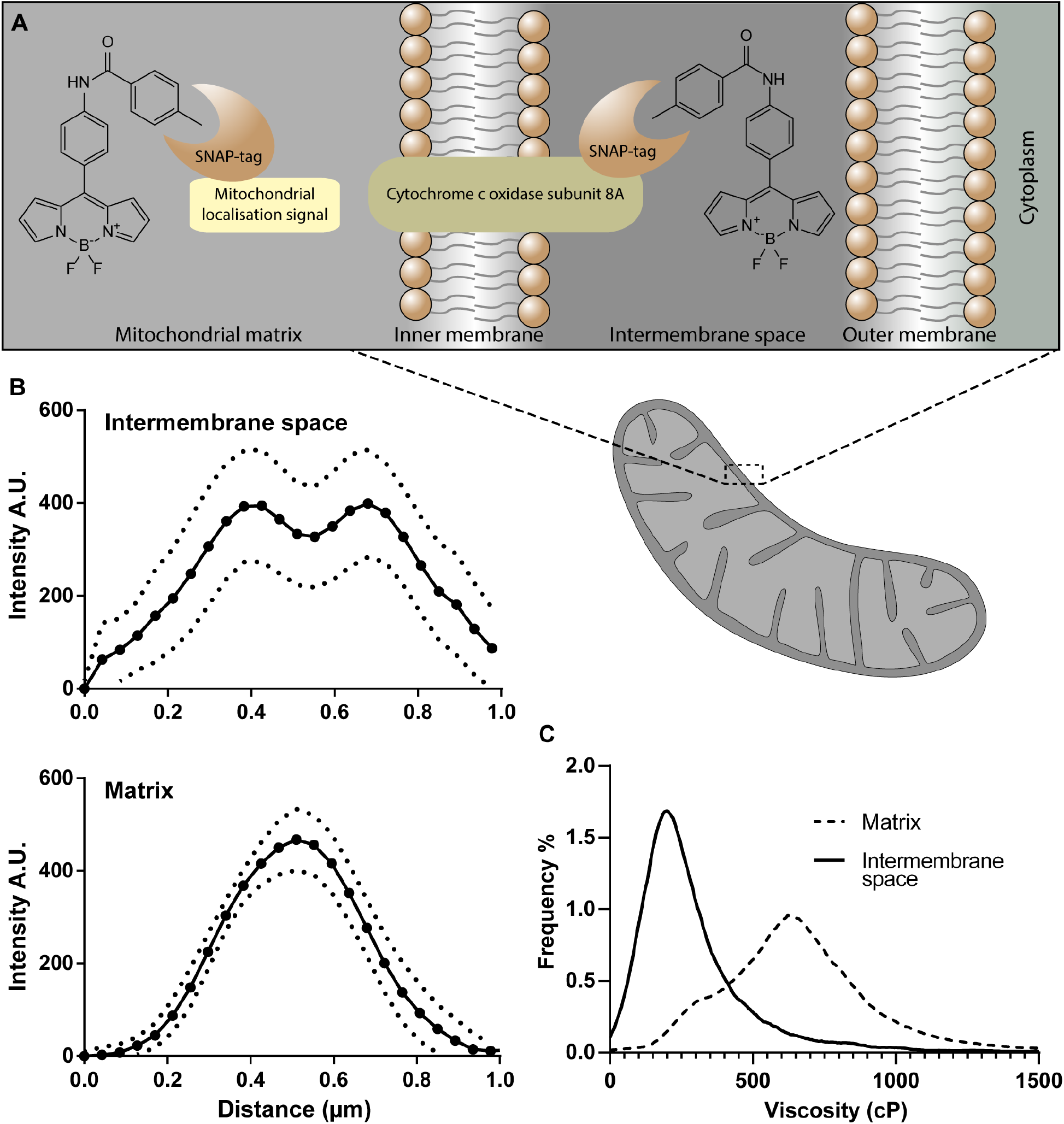
BG-BODIPY reports microviscosity in sub cellular compartments. **A.** Diagram of BG-BODIPY localisation in the mitochondrial matrix and mitochondrial intermembrane space. **B.** Intensity profiles orthogonal to mitochondrial edges showing the single and double peaks of intensity confirming the correct localisation of BG-BODIPY (solid line is the mean of 3 line-scans, dotted lines represent standard deviation). **C.** Histogram of the viscosities reported from FLIM analysis of BG-BODIPY bound to the Cox8/SNAP-tag fusion protein or in the mitochondrial matrix. Mean viscosity for intermembrane space and matrix are respectively 185 cP and 615 cP. A Mann-Whitney test rejects the hypothesis that the two samples have the same median with a p<0.0001.

**Table 1:**
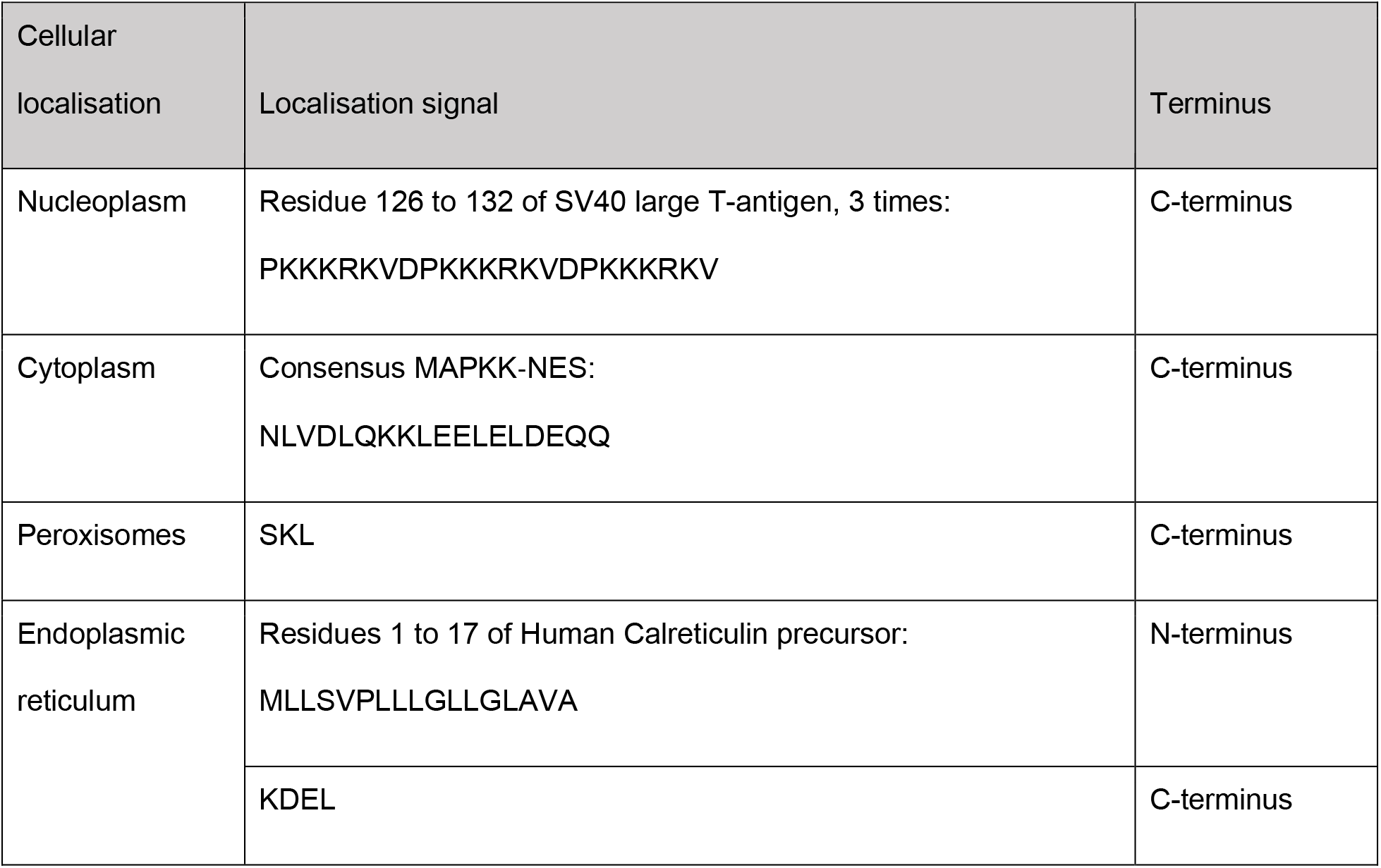
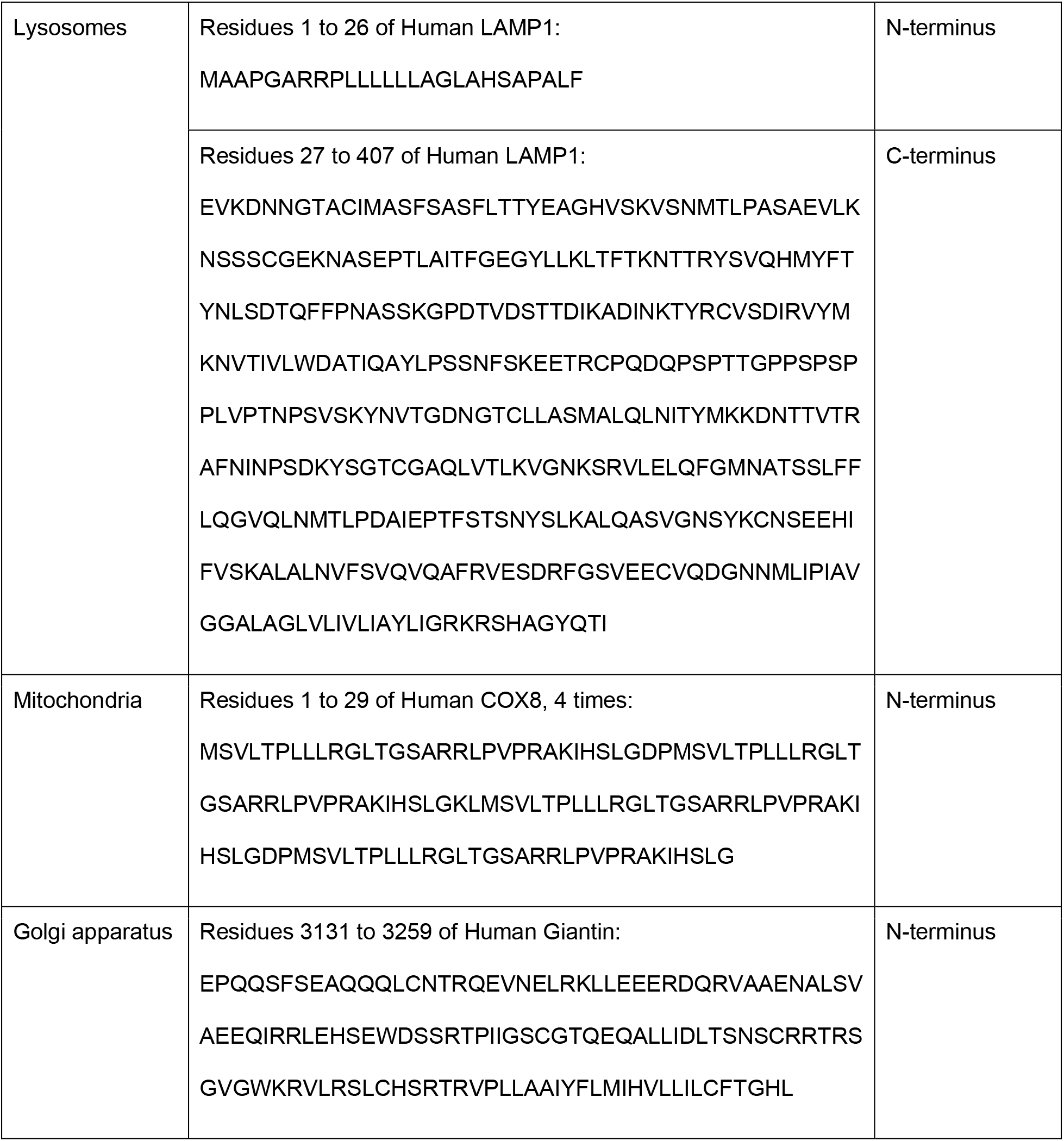
Cellular localisation and respective localisation signals.

### BG-BODIPY can be targeted to measure viscosity in membraneless organelles

To test whether BG-BODIPY could be used to probe cellular compartments formed through liquid-liquid phase separation, we expressed in HeLa cells a construct containing SNAP-tag fused to a C-terminal fragment of the TAR (Trans-activation response element) DNA-binding protein 43 (TDP-43) that lacks the RRM (RNA recognition motifs) domains and is not itself prone to aggregation in physiological circumstances^27^. TDP-43 is a constituent protein of the ubiquitinated inclusions observed in Amyotrophic Lateral Sclerosis (ALS) and frontotemporal dementia, and has been implicated in a number of other dementias, including Alzheimer’s disease^28,29^. Stress granules are cytoplasmic inclusions that sequester actively-translating mRNAs and distinct RNA binding proteins (RBPs) in conditions of oxidative stress^30^. TDP-43 has been shown to associate with stress granules in a host of pathological conditions^31,32^. ZnCl_2_ has been shown to induce the formation of TDP-43 inclusions in the cytoplasm, and has been used to model pathological aggregation of the protein^33^. Therefore, we used ZnCl_2_ to induce stress granules in HeLa cells expressing the TDP-43/SNAP-tag fusion protein^34^.

When the lifetime of BG-BODIPY in the nucleus was measured by FLIM, we found the mean viscosity to be 185 cP (Figure 9C). When cells were treated with 100 µM ZnCl_2_ overnight, a treatment shown robustly to elicit TDP-43 aggregation^35^, we observed distinct populations of foci in the cytoplasm, demonstrating that the truncated TDP-43 is sequestered to stress granules upon oxidative stress (Figure 9A and B). There was wide variation in the viscosities reported from within the stress granules, however we found that the mean viscosity was ~520 cP, a significant increase from TDP-43’s nucleoplasmic microviscous environment.

**Figure 9:**
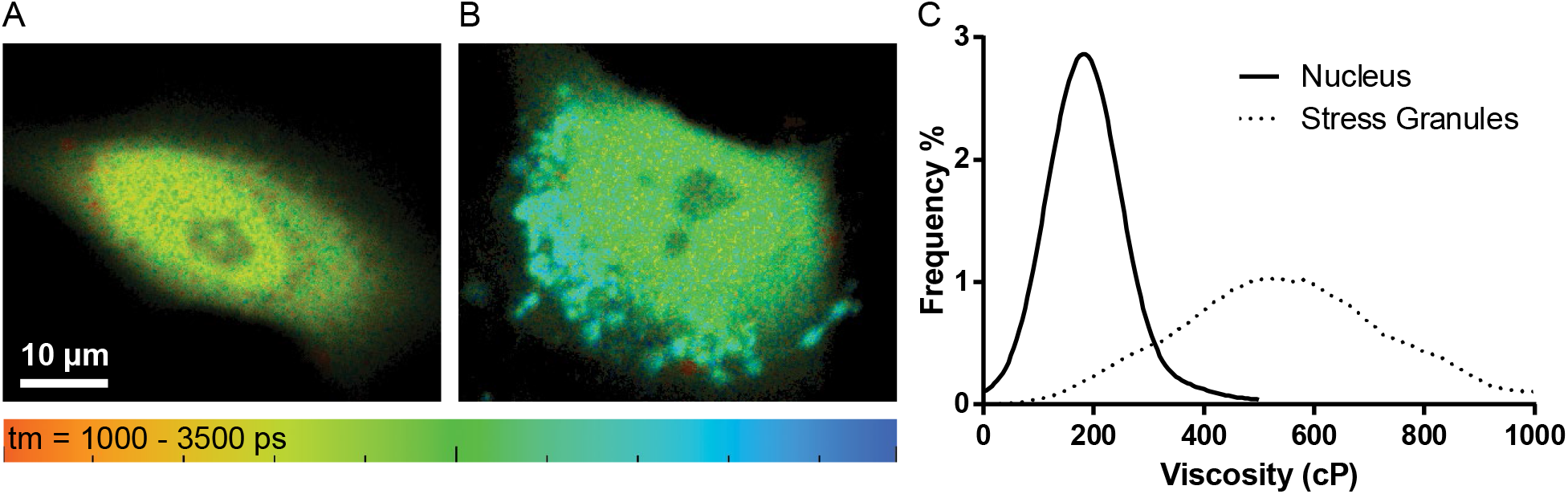
FLIM demonstrates that BG-BODIPY targets to and measures viscosity in membraneless organelles. **A**. C-terminal fragment of TDP43 labelled with BG-BODIPY targets to the nucleoplasm plus a diffuse cytoplasmic signal. **B**. In the presence of ZnCl_2_ the SNAP-tag fusion protein containing the C-terminal fragment of TDP43 forms cytoplasmic aggregates. **C**. Histogram of the viscosities reported from FLIM analysis of BG-BODIPY bound to the TDP43/SNAP-tag fusion protein resting or ZnCl_2_ induced stress granules. Mean viscosity for nuclear TDP43 and stress granules are respectively 185 cP and 520 cP. A Mann-Whitney test rejects the hypothesis that the two samples have the same median with a p<0.0001.

## Discussion

In this study, we demonstrated the usefulness of complementary techniques for measuring viscosity of discrete subcellular locations in living cells. Whilst numerous previous attempts to target molecular rotors in living cells have relied on exploiting the unique properties of given organelles^7,8^, our work represents one of the first attempts at producing a generic mechanism to target BODIPY to any selected organelle or structure. We measured the relative viscosities of a host of organelles, as well as showcasing the BG-BODIPY’s prospects for measuring absolute viscosity in both membrane-bound and membraneless organelles. We did so by CRISPR-Cas9 engineering the *AAVS1* genomic locus in human cells to express the mCherry-SNAP-tag fusion protein targeted to multiple subcellular locations.

FLIM is a powerful technique that provides measurements of molecular rotor fluorescence lifetime, and thus absolute microviscosity, that are independent of probe concentration, but it has several practical limitations. Acquisition of FLIM images occurs over the order of minutes, with weaker signals requiring longer acquisition periods. This precludes the use of FLIM for capturing fast changes in viscosity. Moreover, FLIM requires specialised, expensive equipment not available to most labs. FLIM may be carried out at conventional confocal microscopy resolution, or with the enhancement of two photon imaging. Spatial resolution may be improved by selective targeting of the reporter via expression of organelle-specific fusion proteins bearing the SNAP-tag enzyme. In contrast, using BG-BODIPY with the ratiometric CRISPR cell lines allows us to visualise relative viscosity changes at higher temporal resolution using standard confocal microscopes, and may offer improved spatial resolution via super-resolution methods. Thus, these two approaches represent complementary techniques that can be added to researchers’ toolkits, each for use in appropriate situations.

By using BG-BODIPY in these two methods, we were able to demonstrate its usefulness in probing the microviscous environments of the cell. We showed that there is significant variation in both the mean and the range of the viscosities seen within various cellular organelles. Of particular note is the change as one moves along the secretory pathway. Not only does the mean ratio between BG-BODIPY and mCherry increase from the ER to the Golgi apparatus, but there is also a progressively wider range of viscosities reported as one proceeds along the secretory pathway. We interpret this to mean that the microviscous environment becomes more varied later in the secretory pathway. Whilst the cause of this viscosity variation within an organelle later in the secretory pathway is unclear, it is tempting to consider whether this may be related to the formation of subdomains within later secretory organelles that facilitate separation of cargo with different destinations (secretory granule, constitutive secretion, shuttle vesicle to endocytic pathway).

Our measured organelle viscosity ratios were comparable with those of Chambers et al. (2018)^9^, with the mitochondria having significantly higher mean viscosity than all organelles other than the Golgi apparatus, which was not measured in their study. Using FLIM we were able to determine that the absolute viscosity within the mitochondrial intermembrane space was 185 cP. Furthermore, we were able to distinguish the viscosity in the intermembrane space from that of the mitochondrial matrix by expressing fusion proteins that delivered the SNAP-tag enzyme specifically to one side or the other of the inner mitochondrial membrane. The inner mitochondrial membrane is rich in the unusual lipid cardiolipin^36^ and poses a very stringent barrier, so it is not surprising that the extremely protein-rich matrix space should maintain a higher viscosity despite being only 7nm from the intermembrane space.

Our viscosity ratios in the cytoplasm and nucleoplasm were less concordant with the values reported by Chambers et al, as our probe reported a lower viscosity in the nucleoplasm than the cytoplasm, which may be due to cell type differences. Furthermore, the volume of distribution of the probes may not be comparable since previous attempts have relied upon an untargeted probe that was allowed to spread throughout the cell, whereas our probe was targeted to the nucleus or cytoplasm via either an NLS or an NES, respectively.

Beyond membrane-bound organelles, we demonstrate the possibility of microviscosity measurement within membraneless organelles by targeting BG-BODIPY to proteins involved in forming these organelles. We chose to examine TDP-43 positive stress granules because they represent a pathologically relevant instance of liquid-liquid phase separation. It is thought that the concentration of TDP-43 into stress granules can lead to aggregation of the protein, as seen in numerous neurodegenerative diseases^31^. The ability to measure the dynamic microviscosity environment of TDP-43 in a variety of physiological and disease conditions could lead to important discoveries in prevention of its cellular toxicity. What is more, by applying this technique to proteins involved in other membraneless organelles, we could gain key insight into the more general question of organisation of cellular space through phase separation.

In summary, the techniques presented here represent some powerful new tools for studying live cells. The CRISPR cell lines present a novel means of targeting SNAP-tag compatible probes to various cellular locations, without the disadvantages of transfection and transient overexpression. Any membrane-permeant probe compatible with SNAP-tag is compatible, and these cell lines could be just as easily used with Ca^2+^ or pH-sensing probes as they have been with BG-BODIPY.

We have used both these cell lines, as well as transient transfection with SNAP-tag compatible constructs to demonstrate the potential for FLIM and ratiometric imaging of BG-BODIPY for the measurement of viscosity in living cells. While ratiometric imaging can be useful for relative measurements, it is important to note that there can be fluorescence resonance energy transfer (FRET) and that the calibration curve used to calculate viscosity may not apply to all compartments within the cell. Therefore, the ratiometric method should ideally be used only when FLIM is not available or practical, and only for relative measurements, such as changes across treatments within a single compartment.

Alternatively, using FLIM alone can provide more accurate and reliable viscosity measurements, especially when mCherry or other fluorophores that may act as acceptors for FRET are not present. These techniques allow a single probe to be adapted for use in a variety of circumstances, and facilitate wide-ranging future work in this area. The results presented here represent a small sample of the potential utility of the method, pose many questions about the role of viscosity in cellular processes, and facilitate wide-ranging future work in this area.

## Supporting information

Pytowski et al Movie 1

Pytowski et al Supplementary material

## Abbreviations

BG-BODIPY: benzyl-guanine boron-dipyrromethene
FLIM: fluorescence lifetime imaging microscopy
TDP: TAR DNA-binding protein 43

## Acknowledgments

We gratefully acknowledge the assistance of Cristian Soitu for microfluidic chamber preparation, and Errin Johnson and Raman Dhaliwal from the Dunn School EM Facility for TEM sample preparation, labelling and imaging. We thank the Micron Oxford Advanced Bioimaging Unit (funded from Wellcome Trust Strategic Award 107457/Z/15/Z) for technical assistance, discussions and advice, and Michal Maj from the Flow Cytometry Facility - Sir William Dunn School of Pathology for cell sorting.

## Author contributions

L.P., A.C.F. and D.J.V. conceived and planned the experiments. A.C.F. carried out the FLIM experiments. L.P. carried out the other light microscopy experiments. L.P. and Z. E. H. created the CRISPR’d cell lines. N.M. synthesized the BG-BODIPY under T.D. direction. L.P, A.C.F. and D.J.V contributed to the interpretation of the results. A.C.F. took the lead in writing the initial manuscript with support from L.P. and D.J.V. Data visualisation by L.P. All authors provided critical feedback and helped shape the research, analysis and manuscript.

## Competing financial interests

Authors declare no conflicts of interest with the contents of this article.

## Notes

### Competing Interest Statement

The authors have declared no competing interest.

### Summary of Updates

This revision reports a substantial development of the image analysis pipeline for FLIM and ratiometric images, including the use of single and double Gaussian fits to the data, tested for quality via examination of SSD (sum of squared differences).

