## Supplementary material for "Mapping viscosity in discrete subcellular locations with a BODIPY based fluorescent probe": Pytowski et al Supplementary material

**Supplementary figures**

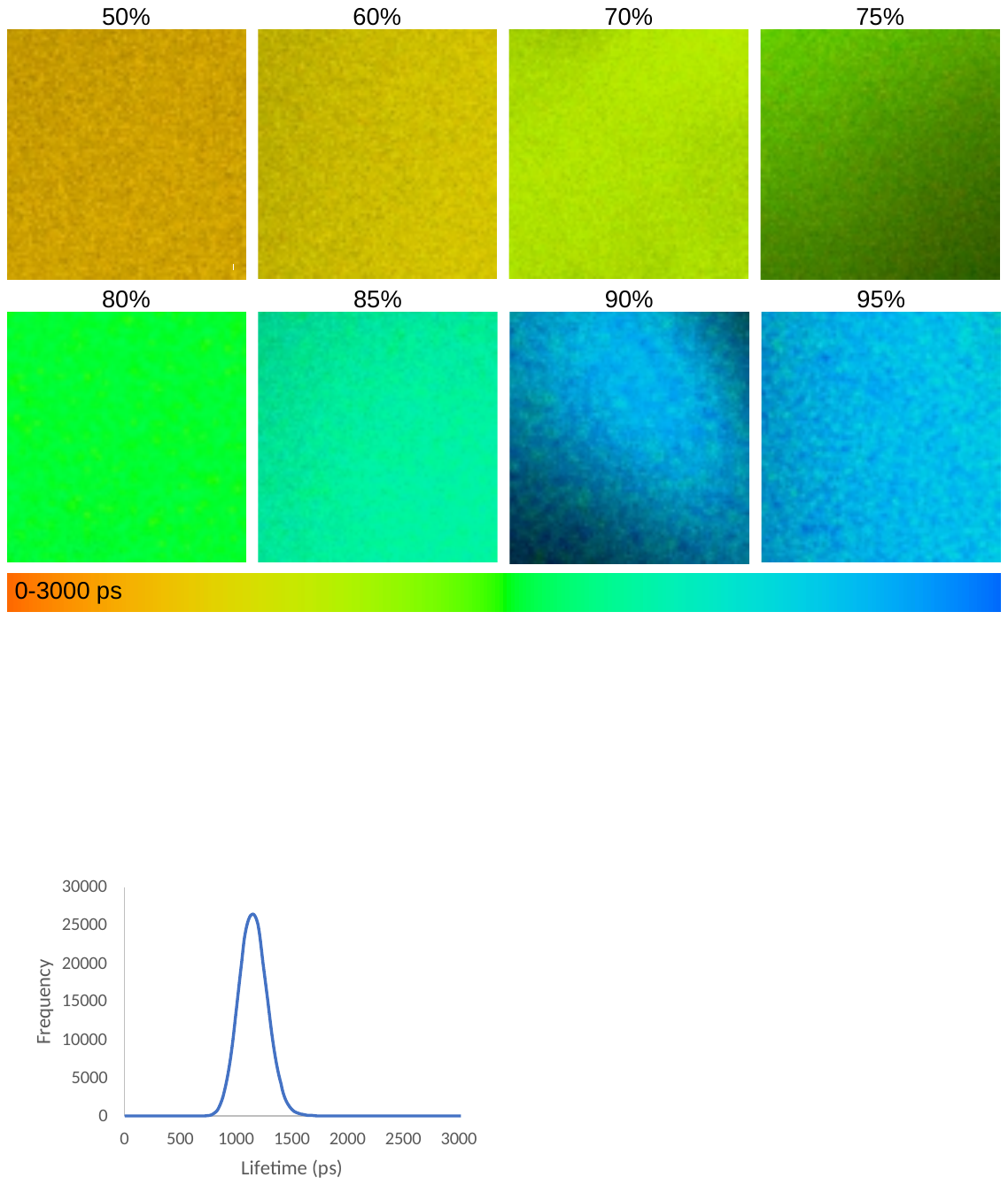

**Supplementary figure 1: Sample images used for calibration curve. A.** Sample images used for calibration curve. No aggregation of BG-BODIPY was observed. Percentages are the glycerol content. **B.** Output of SPC-Image for a representative 75% glycerol sample, showing the time distribution histogram of individual pixel counts per time bin (blue dots) and the resulting curve fit (red line). **C**. Lifetime frequency histogram from data in (B) showing single predominant lifetime.

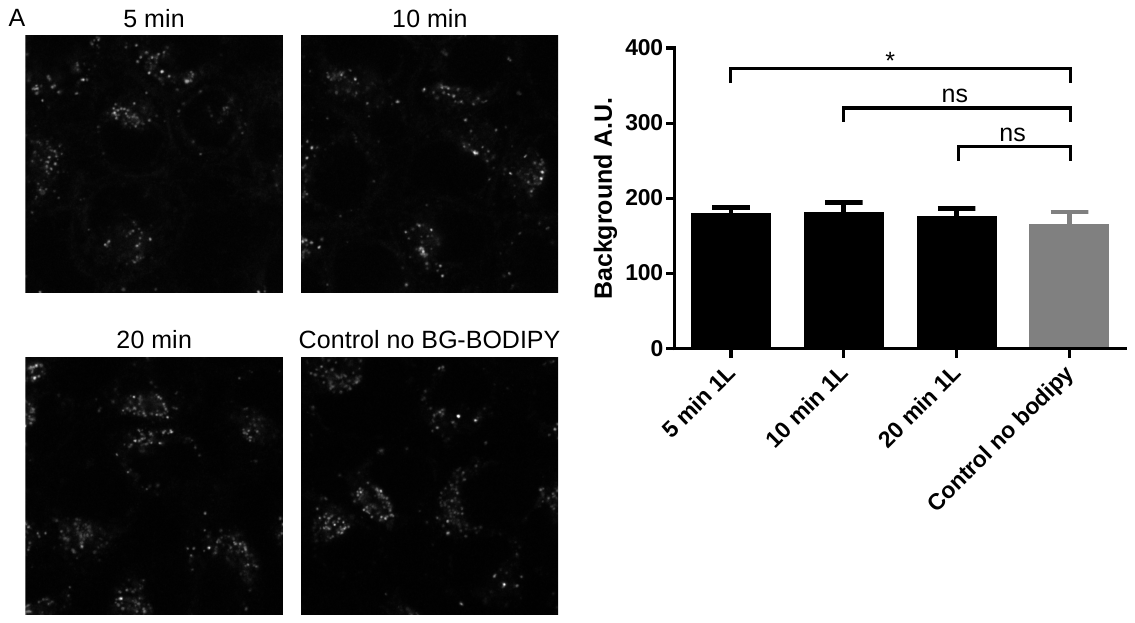

**Supplementary figure 2:** **Non-specific background of unbound BG-BODIPY can be removed by washing with DMEM containing fatty acid-free BSA . A**. HeLa cells containing no SNAP-Tag were imaged after incubation with BG-BODIPY to mimic the usual labelling protocol, followed by three 5% fatty acid free BSA washes for 5, 10 or 20 minutes. The results are compared to a sample of HeLa cells that had not been exposed to BG-BODIPY imaged under the same conditions **B**. Quantification of the residual abundance of BG-BODIPY after the washes.

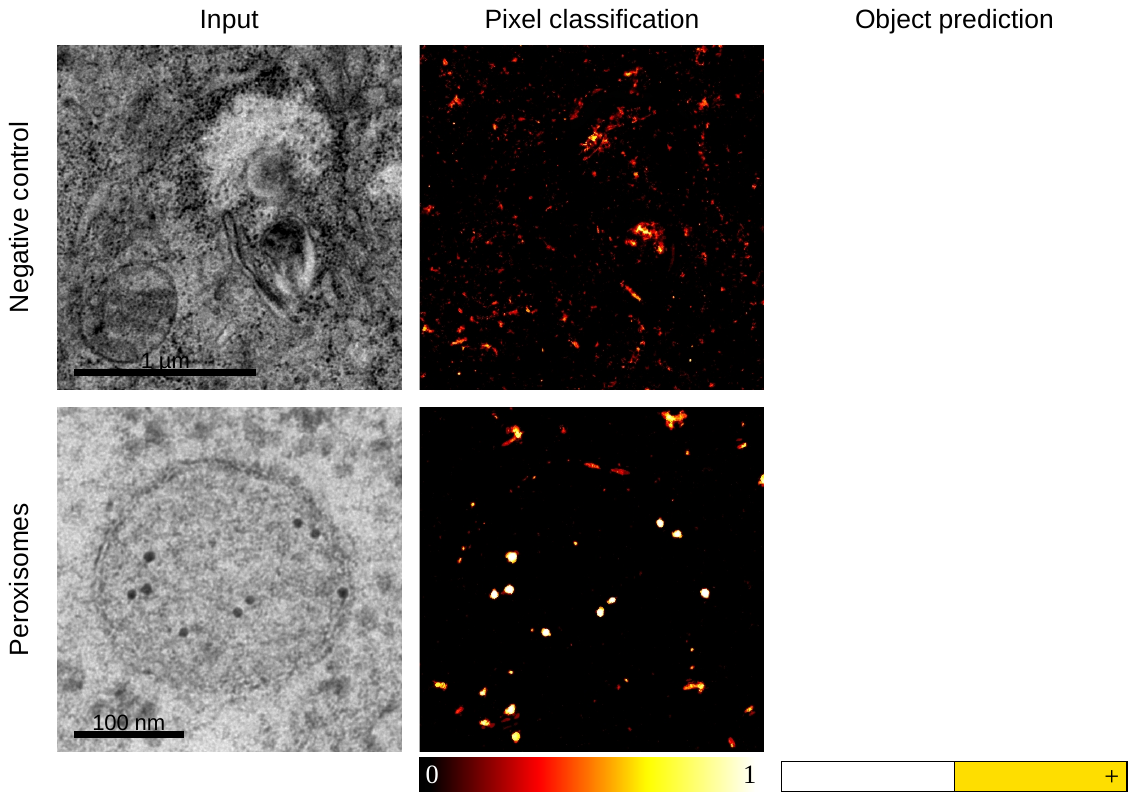

**Supplementary figure 3:** Identification of gold particles in immunolabelled images by automatic segmentation. Gold particles were segmented using Ilastik. Pixels were scored from 0 to 1, 1 being the highest likelihood of being a gold particle. Object were predicted based on multiple parameters including shape and size. Yellow objects are gold particles, white objects are not gold particles.

**Supplementary Methods**

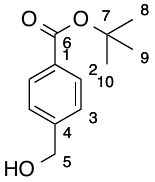
**Synthesis of** ***tert*-butyl 4-(hydroxymethyl)benzoate (Compound 1)**

In a flame dried round bottom flask, equipped with a stirrer bar, 4-formyl benzoic acid (3.00 g, 20.0 mmol) was dissolved in *tert*-butanol (300 mL) and degassed with argon for 10 minutes. Di-*tert*-butyl dicarbonate (6.89 mL, 30.0 mmol) was then added quickly followed by DMAP (0.244 g, 2.00 mmol). The reaction mixture was heated to 35 °C and left to stir for 16 h. Additional di-*tert*-butyl dicarbonate (4.60 mL, 20.0 mmol) was then added and the reaction left for a further 24 h. Excess solvent was then removed on a rotatory evaporator and left to dry *in vacuo*. This crude product was then stirred in dry methanol (80 mL) until dissolved. NaBH_4_ (0.907 g, 24.0 mmol) was added in five batches over 30 minutes. The reaction was left to stir for 18 h before the addition of additional NaBH_4_ (150 mg, 3.97 mmol) and the reaction left to stir for a further 2 h. The solvent was then removed on the rotatory evaporator before the addition of EtOAc (400 mL). The organic layer was then washed with brine (300 mL) and NaHCO_3_ (300 ml), dried over Na_2_SO_4_ and the solvent removed under vacuum. This crude product was purified *via* flash column chromatography to yield the desired product (2.69 g, 12.9 mmol, 64.6%) as a white solid.

**R_f_** = 0.27 (Pentane : Et_2_O; 1 : 1); **M.p.** = 32 °C; **^1^H NMR** (400 MHz, CDCl_3_) δ 7.93 (d, *J* = 7 .8 Hz, 2H, C*H(2)*), 7.38 (d, *J* = 7.8 Hz, 2H, C*H(3)*), 4.72 (s, 2H, C*H_2_(5)* ), 1.58 (s, 9H, C*H_3_(8-10)*); **^13^C NMR** (101 MHz, CDCl_3_) δ 166.2 (*C(6)*), 146.0 (*C(4)*), 131.5 (*C(1)*), 130.1 (*C(2)*), 126.7(*C(3)*), 81.5 (*C(7)*), 65.1 (*C(5)*), 28.6 (*C(8-10)*); **FTIR** ν_max_ (neat) 3423 (w), 2961 (w), 1711 (s), 1673 (m), 1365 (s), 1288 (s), 1153 (s), 1115 (s), 1019 (m), 827 (m), 750 (m)**; HRMS** (ESI+) calculated for [C_12_H_16_O_3_Na]^+^: 231.0992. Found: 231.0993 (Δ = 0.53 ppm).

Analytical data matches that in the literature^1^*.*

**Synthesis of** **1-(2-amino-9*H*-purin-6-yl)-1-methylpyrrolidin-1-ium chloride (Compound 2)**

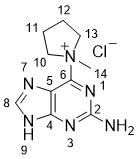

In a flame dried round bottom flask, 6-chloro-guanine (8.00 g, 47.2 mmol) was dissolved in DMF (320 mL) and heated to 40 °C until the solid was dissolved. After cooling to room temperature, 1-methyl-pyrrolidine (11.3 mL, 85.2 mmol) was added and the reaction was left to stir for 27 hours. The white precipitate was filtered, washed with diethyl ether (100 mL) and dried under vacuum to yield the desired product in 41.7% yield (5.01 g, 19.7 mmol) as a white solid.

**M.p.** 194 °C; **^1^H NMR** (400 MHz, Methanol-*d*_4_) δ 8.26 (s, 1H, C*H(8)*), 4.96 – 4.84 (m, 2H, C*H_2_(10*), 4.20 – 3.95 (m, 2H, C*H_2_(13)*), 3.81 (s, 3H, C*H_3_(14)*), 2.57 – 2.36 (m, 2H, C*H_2_(11)*), 2.38 – 2.02 (m, 2H, C*H_2_(12)*); **^13^C NMR** (126 MHz, Methanol-*d*_4_) δ 161.1 (*C(2)*), 160.4 (*C(4)*), 153.0 (*C(6)*), 144.0 (*C(8)*), 117.7 (*C(5)*), 65.8 (*C(10,13)*), 52.7 (*C(14)*), 22.8 (*C(11,12)*); **FTIR** ν_max_ (neat) 3346 (w), 3290 (w), 3181 (w), 2979 (w), 2937 (w), 2893 (w), 2756 (m), 1624 (m), 1564 (m), 1295 (m), 921 (m), 620 (s) cm^1^; **HRMS** (ESI-) calculated for [C_10_H_14_N_6_Cl]^+^: 253.0974. Found: 253.0972 (Δ = −0.96 ppm).

Analytical data matches that in the literature^2^.

**Synthesis of** ***tert*-butyl 4-(((2-amino-9*H*-purin-6-yl)oxy)methyl)benzoate (Compound 3)**

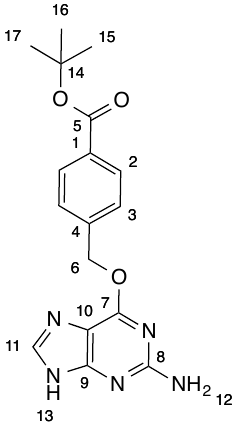

In a flame dried round bottom flask *tert*-butyl 4-(hydroxymethyl)benzoate was dissolved in dry DMF (7 mL) and degassed with argon for 10 minutes. NaH (0.418 g, 10.5 mmol, 60% in mineral oil) was then added in 3 batches over 15 minutes. 4-(Dimethylamino)pyridine (43.0 mg, 0.349 mmol) and 1-(2-amino-9*H*-purin-6-yl)-1-methylpyrrolidin-1-ium chloride (0.888 g, 3.49 mmol) were then quickly added and the reaction was left to stir for 26 h. Water (1.00 mL) was then added slowly followed by EtOAc (30 mL). The organic layer was washed with brine (6 x 30 mL) before being dried over Na_2_SO_4_ and the excess solvent removed under vacuum. The crude product was purified *via* flash column chromatography (5 % MeOH / DCM) to yield the desired product (0.150 g, 0.439 mmol, 12.6%) as a white solid.

**M.p.** = >350 °C; **R_f_** = 0.40 (MeOH : DCM; 1 : 9); **^1^H NMR** (500 MHz, Acetone-*d*_6_) δ 11.77 (bs, 1H, N*H(13)*), 7.96 (d, *J* = 8.0 Hz, 2H, C*H(2)*), 7.86 (s, 1H, C*H(11)*), 7.61 (d, *J* = 8.0 Hz, 2H, C*H(3)*), 5.81 (bs, 2H, N*H_2_(12)*), 5.60 (s, 2H, C*H_2_(6)*), 1.58 (s, 9H, C*H_3_(15-17)*); **^13^C NMR** (126 MHz, Acetone-*d*_6_) δ 165.8 (*C(5)*), 161.2 (*C(8)*), 160.8 (*C(7)*), 156.5 (*C(9)*), 142.9 (*C(4)*), 138.5 (*C(11)*), 132.4 (*C(1)*), 130.1 (*C(2)*), 128.7 (*C(3)*), 115.2 (*C(10)*), 81.3 (*C(14)*), 67.3 (*C(6)*), 28.3 (*C(15-17)*); **FTIR** ν_max_ (neat) 3484 (w), 2104 (w), 2981 (m), 2359 (w), 1707 (m), 1618 (s), 1580 (s), 1395 (m), 1283 (s), 1162 (m), 1283 (s), 754 (m); **HRMS** (ESI-) calculated for [C_17_H_18_O_3_N_5_]^-^: 340.1415. Found: 340.1412 (Δ = -0.99 ppm).

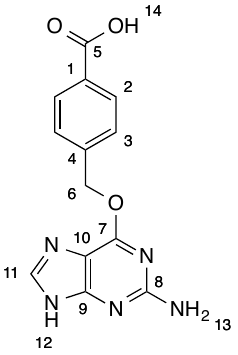
**Synthesis of 4-(((2-amino-9*H*-purin-6-yl)oxy)methyl)benzoic acid (Compound 4)**

*tert*-Butyl 4-(((2-amino-9*H*-purin-6-yl)oxy)methyl)benzoate (0.386 g, 1.13 mmol) was dissolved in a flame dried round bottom flask, equipped with a stirrer bar, in dry dichloromethane (10 mL). TFA (1.12 mL, 14.7 mmol) was then added and the reaction was left to stir for 4.5 h. Dichloromethane was removed under vacuum to yield a white solid which was washed with EtOAc (3 x 50 mL). The solid was dried under vacuum to yield the TFA salt of the desired product (0.340 g, 0.852 mmol, 75.3%) as a white solid.

**M.p.** = >350 °C; **^1^H NMR** (500 MHz, DMSO) δ 13.01 (bs, 1H, N*H(11)*) 8.34 (s, 1H, *CH(11)*), 7.98 (d, *J* = 8.4 Hz, 2H, C*H(2)*), 7.64 (d, *J* = 8.4 Hz, 2H, C*H(3)*), 7.10 (bs, 2H, N*H2(13)*), 5.63 (s, 2H, C*H2(6)*); **^13^C NMR** (126 MHz, DMSO) δ 167.0 (*C(5)*), 158.8 (*C(7)*), 158.6 (*C(8)*), 154.2 (*C(9)*), 140.9 (*C(4)*), 140.3 (*C(11)*), 130.5 (*C(1)*), 129.4 (*C(2)*), 128.1 (*C(3)*), 108.8 (*C(10)*), 67.0 (*C(6)*); **FTIR** νmax (neat) 3514 (w), 3136 (m), 2607 (m), 1638 (m), 1623 (s), 1590 (s), 1429 (m), 1280 (s), 1129 (s), 750 (s); **HRMS** (ESI+) calculated for [C_13_H_12_O_3_N_5_]+: 286.0935. Found: 286.0936 (Δ = 0.46 ppm).

**Synthesis** **of *N,N*´-difluoroboryl-5-(4-aminophenyl)-dipyrromethene (Compound 5)**
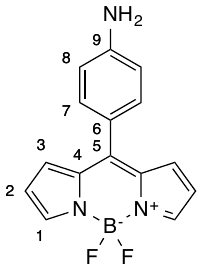

**General Procedure:**

**Step I:** Pyrrole (100.0 equiv.) and aldehyde (1.00 equiv.) were added to a flame dried round bottom flask, equipped with a stirrer bar and degassed with argon for 30 – 60 minutes. A catalytic amount of InCl_3_ (0.10 equiv.) was then added and the solution was stirred under argon at room temperature for stated time. NaOH (3.00 equiv.) was then added and the reaction mixture left to stir for a further 45 minutes. The reaction mixture was filtered under vacuum before excess pyrrole was removed on a rotatory evaporator equipped with a dry ice, acetone bath cooled collection trap. The resultant crude product was then carried through to the next step.

**Step II:** To a flame dried round bottom flask, the crude product from step I was dissolved in toluene (20.0 mL/mmol) and stirred under argon for 30 - 60 minutes. 2,3-Dichloro-5,6-dicyano-1,4-benzoquinone (1.00 equiv.) was then added and the reaction mixture left to stir for 30 - 60 minutes in the dark. Et_3_N (7.00 equiv.) and BF_3_•OEt_2_ (7.00 equiv.) were then added in quick succession before the reaction was left to stir for the indicated time. The reaction mixture was then decanted with toluene (3 x 20 mL/mmol) and washed with water (4 x 50 mL/mmol) before the combined organic layers were dried over Na_2_SO_4_, filtered and the remaining solvent removed under vacuum. The crude product was then purified *via* silica gel column chromatography to give the desired product.

4-Nitrobenzaldehyde (2.00 g, 13.2 mmol) was subjected to the General Procedure above. The reaction was left to stir for 3 h (Step I) then 18 h (Step II) before being purified by flash column chromatography (dichloromethane). The purified product was then dissolved in EtOH (220 mL) in a flame dried round bottom flask, equipped with a stirrer bar, and 10 wt% Pd/C (160 mg) was added. Hydrazine hydrate (2.10 mL, 43.3 mmol) was then added and the reaction was heated at reflux for 2 h. The crude reaction mixture was then filtered through celite, diluted with water (75 mL) and extracted with dichloromethane (5 x 50 mL). The combined organic layers were dried over Na2SO4, filtered and the solvent removed under vacuum. The crude product was then purified by flash column chromatography to yield the desired product (0.606 g, 2.1406 mmol, 16.2 %) as a red solid.

**Rf** = 0.28 (DCM) [Vis, orange]; **M.p.** = 103 °C; **^1^H NMR** (500 MHz, CDCl3) δ 7.89 (s, 2H, C*H(1)*), 7.44 (d, *J* = 8.5 Hz, 2H, C*H(8)*), 7.02 (d, *J* = 4.1 Hz, 2H, C*H(3)*), 6.77 (d, *J* = 8.5 Hz, 2H, C*H(7)*), 6.54 (dd, *J* = 4.1, 1.8 Hz, 2H, C*H(2)*), 4.11 (s, 2H, N*H2*); **^13^C NMR** (126 MHz, CDCl3) δ 149.8 (*C(9)*), 148.3 (*C(5)*), 142.7 (*C(1)*), 134.7 (*C(4)*), 133.0 (*C(7)*), 131.2 (*C(3)*), 124.0 (*C(6)*), 118.0 (*C(2)*), 114.5 (*C(8)*); **^19^F NMR** (470 MHz, CDCl3) δ -145.07, -145.12, -145.19, -145.26; **FTIR** νmax (neat) 3514 (w), 3297 (w), 3136 (m), 2861 (w), 1638 (m), 1623 (s), 1590 (s), 1429 (m), 1280 (s), 1178 (s), 751 (s); **UV-Vis** (DCM): λmax (nm) = 497 (ε = 7951 cm^-1^M^-1^); **HRMS** (ESI-) calculated for [C_15_H_11_BF_2_N_3_]-: 282.1020. Found: 282.1012 (Δ = -1.3 ppm).

**Synthesis of BG-BODIPY (Compound 6)**

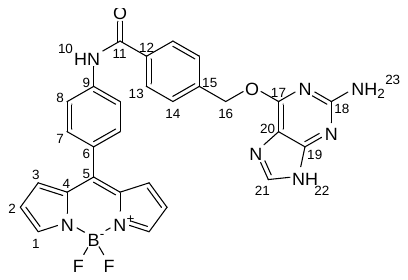

In a flame dried vial, *N,N*´-difluoroboryl-5-(4-aminophenyl)-dipyrromethene (84.9 mg, 0.300 mmol), 4-(((2-amino-9*H*-purin-6-yl)oxy)methyl)benzoic acid (85.6 mg, 0.214 mmol), and *N,N*-diisopropylethylamine (130 μL, 0.450 mmol) were dissolved in dry *N,N*-dimethylformamide (1.50 mL). The reaction mixture was cooled to 0 °C, 1-[bis(dimethylamino)methylene]-1*H*-1,2,3-triazolo[4,5-b] pyridinium 3-oxide hexafluorophosphate (171 mg, 0.450 mmol) was added and the reaction was left to stir for 44 h. The reaction was quenched with NH_4_Cl (aq.) (10 mL) and diluted with dichloromethane (20 mL). The aqueous layer was extracted with dichloromethane (3 x 50 mL) before the combined organic layers were washed with brine (3 x 100 mL), dried over Na_2_SO_4_ and excess solvent removed under vacuum. The crude product was purified via flash column chromatography (3% MeOH: dichloromethane) to yield the desired product (89.4 mg, 0.160 mmol, 54.1%) as an orange solid.

**M.p.** = 188 °C decomposition; **Rf** = 0.23 (MeOH : DCM; 1 : 9); [Vis, yellow]; **^1^H NMR** (500 MHz, DMSO) δ 12.50 (bs, 1H, N*H(22)*), 10.64 (s, 1H, N*H(10)*), 8.12 (s, 2H, C*H(1)*), 8.06 (d, *J* = 8.7 Hz 2H, C*H(13)*), 8.01 (d, *J* = 8.3 Hz 2H, C*H(8)*), 7.87 (s, 1H, C*H(21)*), 7.70 (d, *J* = 8.7 Hz, 2H, C*H(14)*), 7.68 (d*, J* = 8.3 Hz 2H, C*H(7)*) 7.10 (d, *J* = 4.1 Hz 2H, C*H(3)*), 6.70 (dd, *J* = 4.1, 1.7 Hz 2H, C*H(2)*), 6.32 (s, 2H, N*H2(23)*), 5.61 (s, 2H, C*H2(16)*); **^13^C NMR** (126 MHz, DMSO) δ 165.8 (*C(11)*), 159.6 (*C(17)*), 159.4 (*C(18)*), 155.6 (*C(19)*), 146.9 (*C(12)*), 144.2 (*C(1)*), 142.3 (*C(15)*), 140.8 (*C(9)*), 138.3 (*C(21)*), 134.2 (*C(5)*), 134.0 (*C(4)*), 131.7 (*C(14,3)*), 129.3 (*C(6)*), 128.1 (*C(7)*), 128.0 (*C(8)*), 119.8 (*C(13)*), 119.1 (*C(2)*), 113.0 (*C(20)*), 66.1 (*C(16)*); **^19^F NMR** (377 MHz, Methanol-*d*4) δ -146.19, -146.26, -146.34, -146.41; **FTIR** νmax (neat) 3629 (w), 3405 (w), 2360 (w), 1586 (m), 1550 (m), 1411 (s), 1355 (w), 1077 (s), 974 (w), 834 (s); **UV-Vis** (MeOH): λmax (nm) = 497 (ε = 29,389 cm^-1^M^-1^); **HRMS** (ESI+) calculated for [C_28_H_22_BF_2_N_8_O_2_]+: 551.1921. Found: 551.1922 (Δ = 0.12 ppm).

**NMR spectra of synthetic compounds**

***tert*-Butyl 4-(hydroxymethyl)benzoate (Compound 1)**

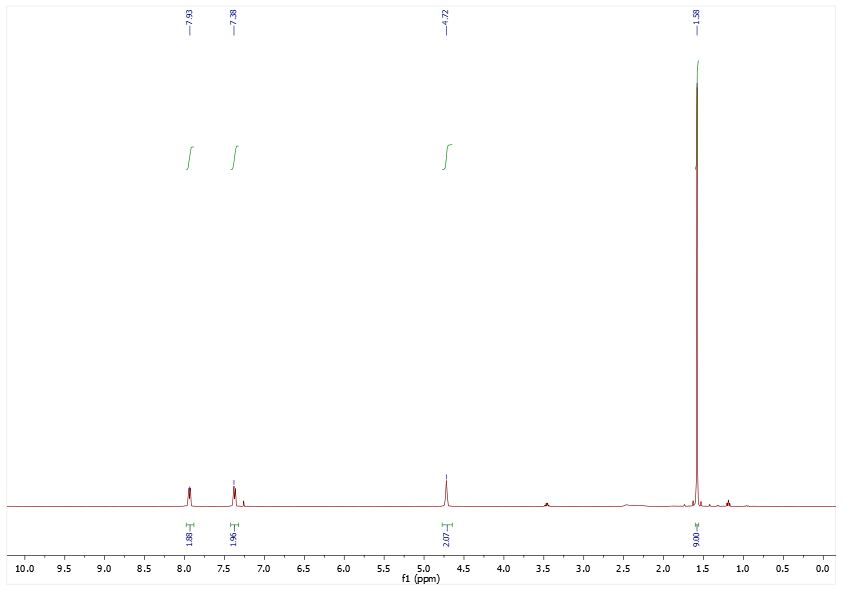
^1^H NMR Spectrum

^13^C NMR Spectrum

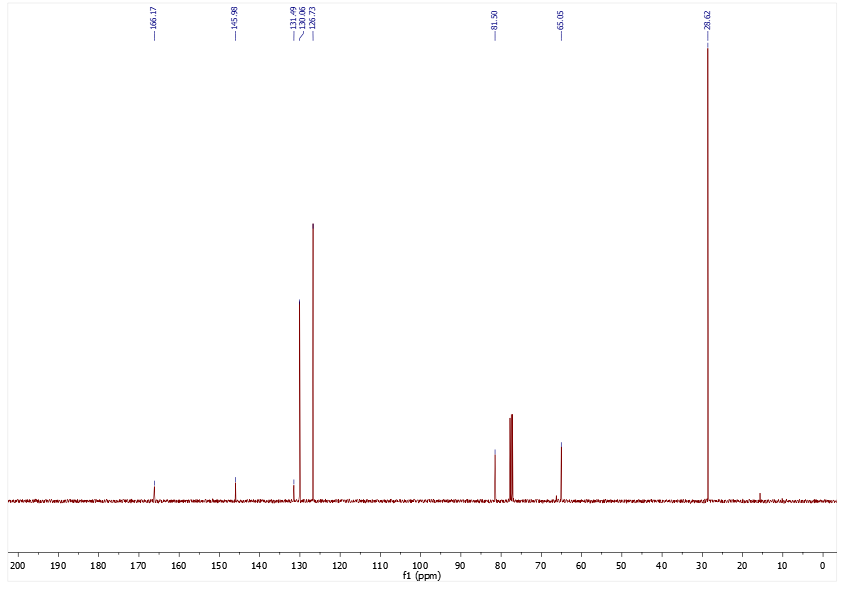

**1-(2-Amino-9*H*-purin-6-yl)-1-methylpyrrolidin-1-ium chloride (Compound 2)**

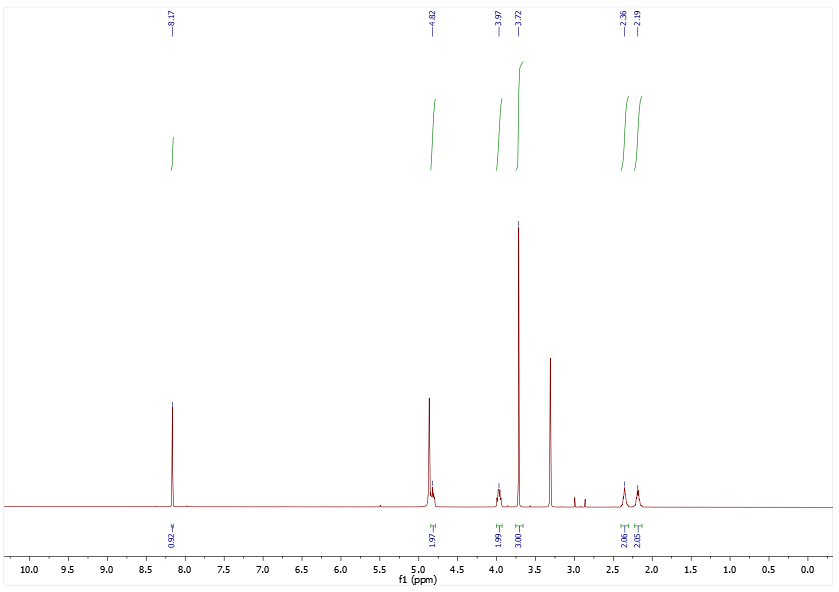
^1^H NMR Spectrum

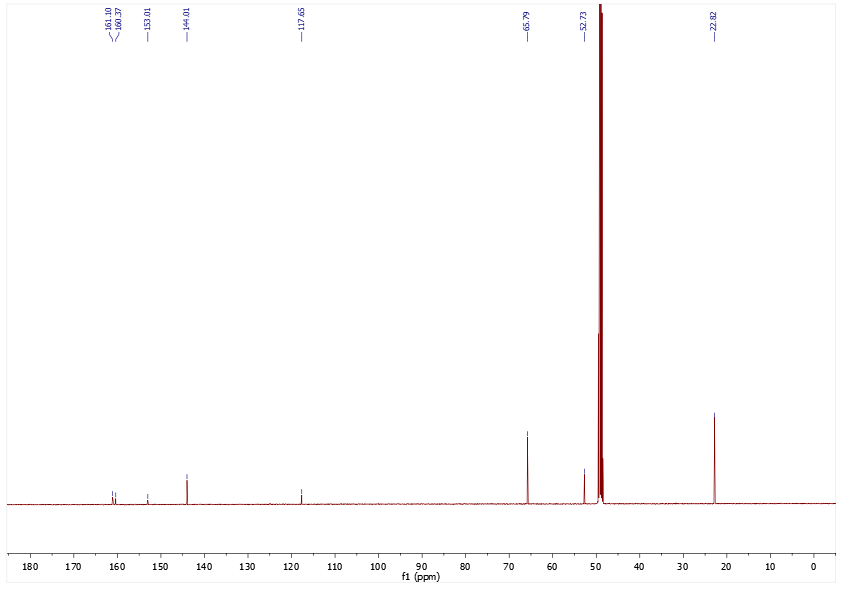
^13^C NMR Spectrum

***tert*-Butyl 4-(((2-amino-9*H*-purin-6-yl)oxy)methyl)benzoate (Compound 3)**

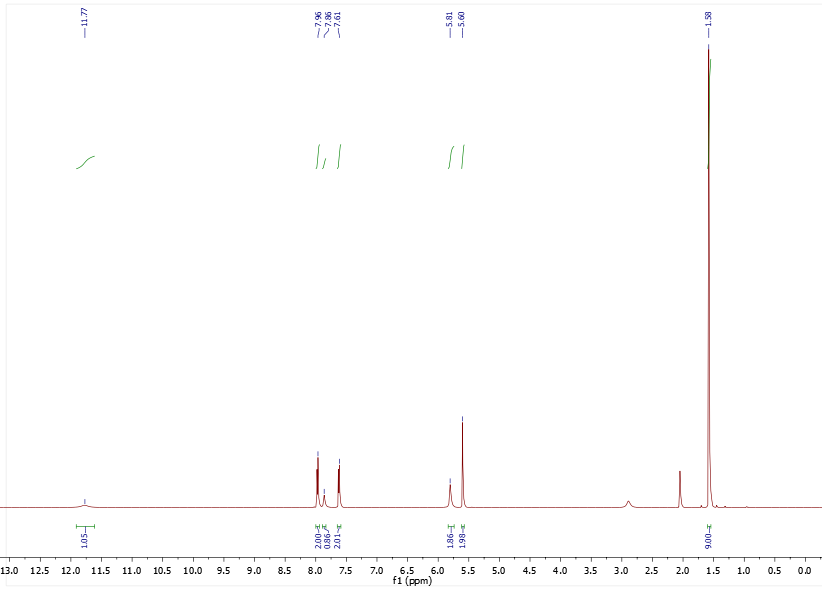
^1^H NMR Spectrum

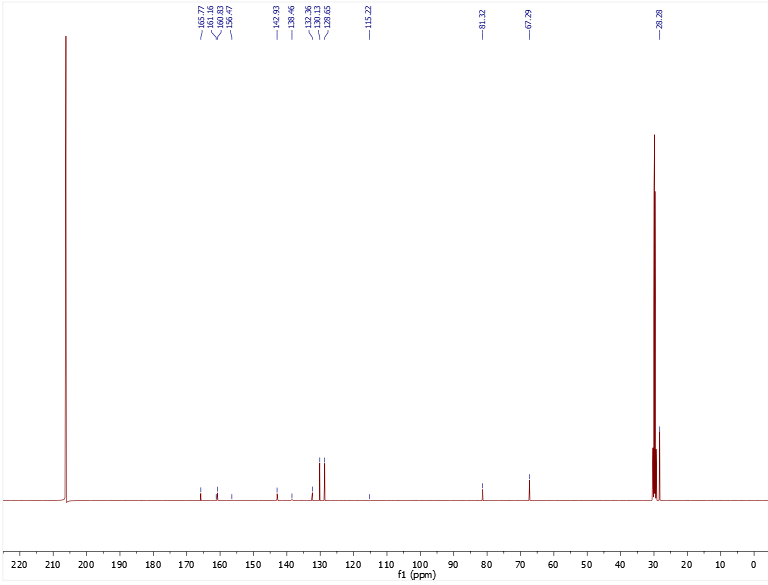
^13^C NMR Spectrum

**4-(((2-Amino-9*H*-purin-6-yl)oxy)methyl)benzoic acid (Compound 4)**

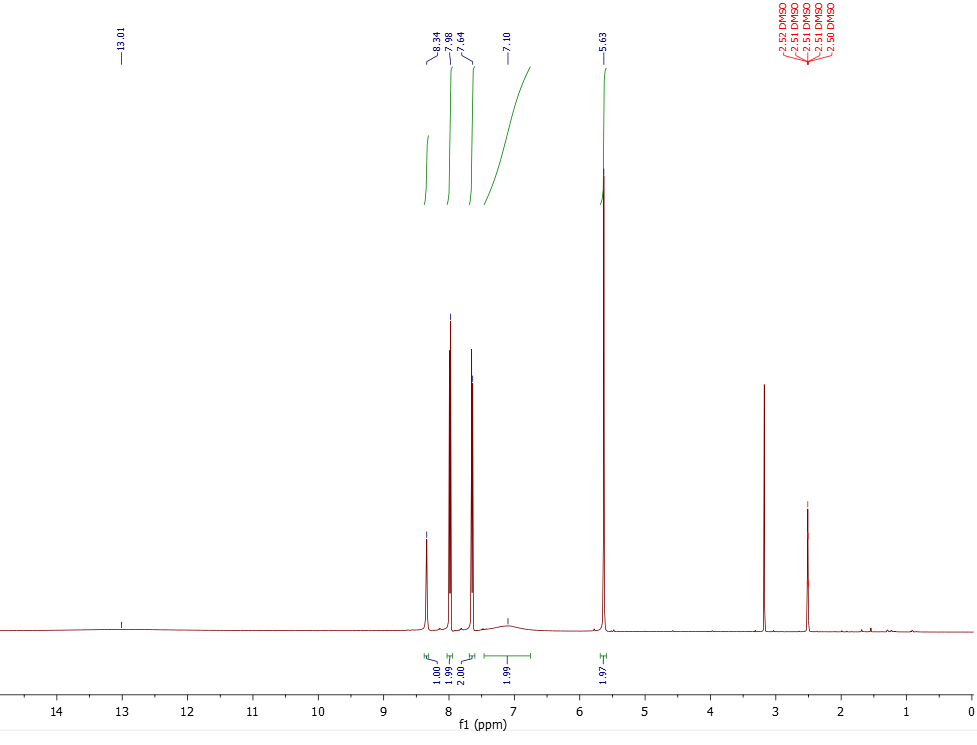
^1^H NMR Spectrum

^13^C NMR Spectrum

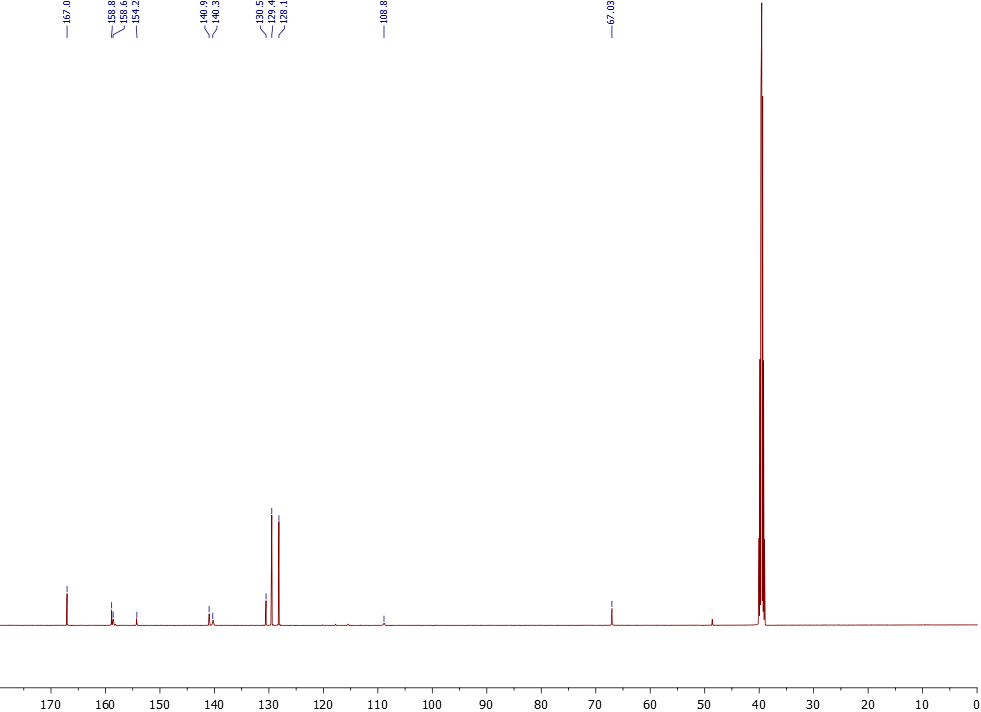

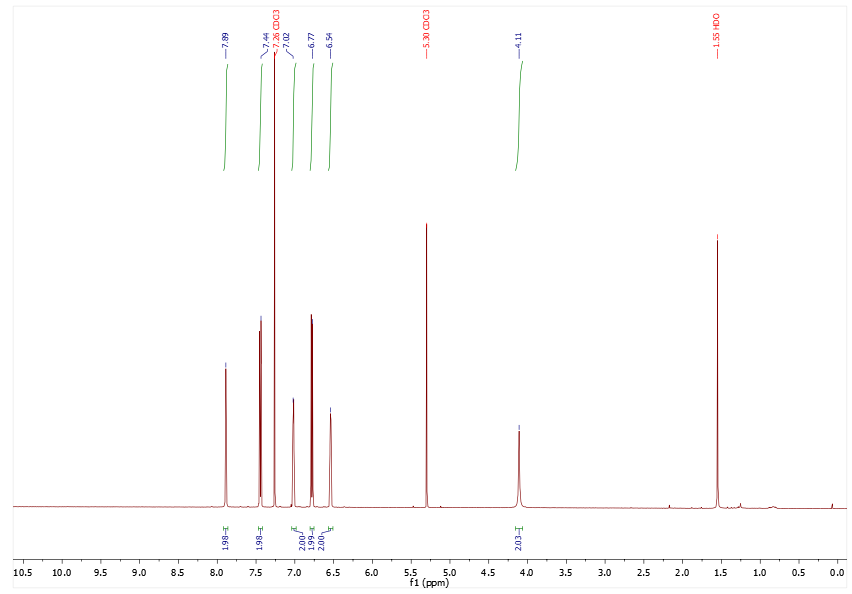
***N,N*´-Difluoroboryl-5-(4-aminophenyl)-dipyrromethene (Compound 5)**

^1^H NMR Spectrum

^13^C NMR Spectrum

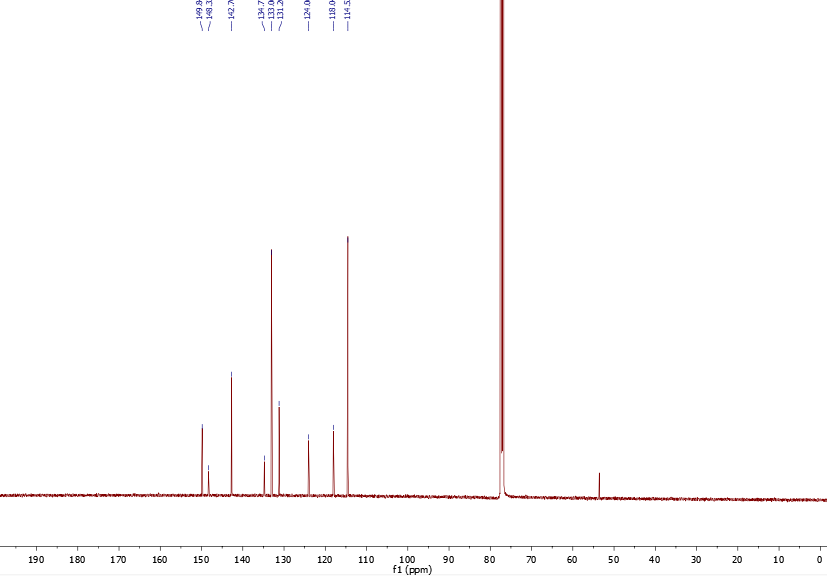

^19^F NMR Spectrum

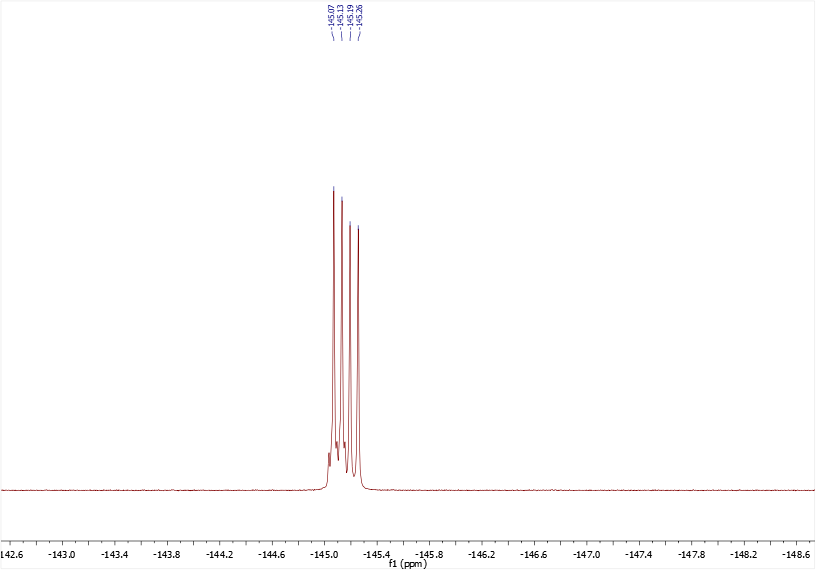

**BG-BODPIY (Compound 6)**

^1^H NMR Spectrum

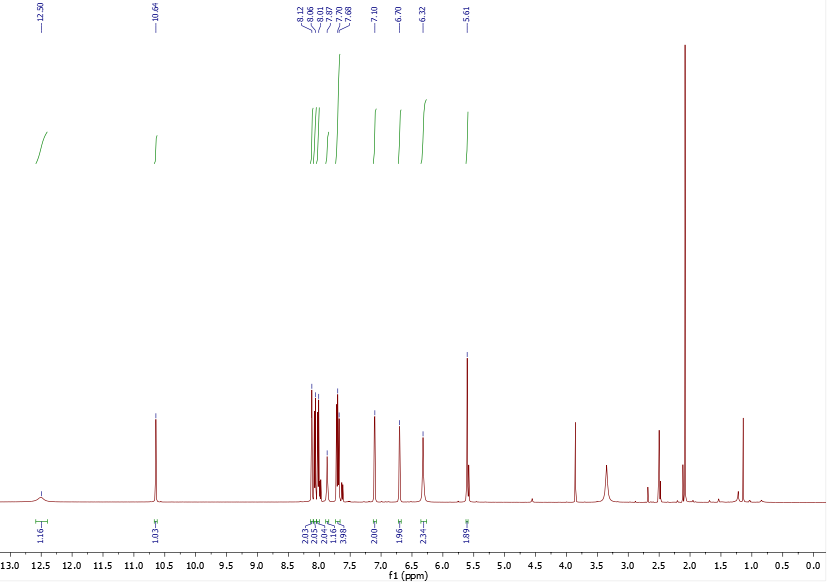

^13^C NMR Spectrum

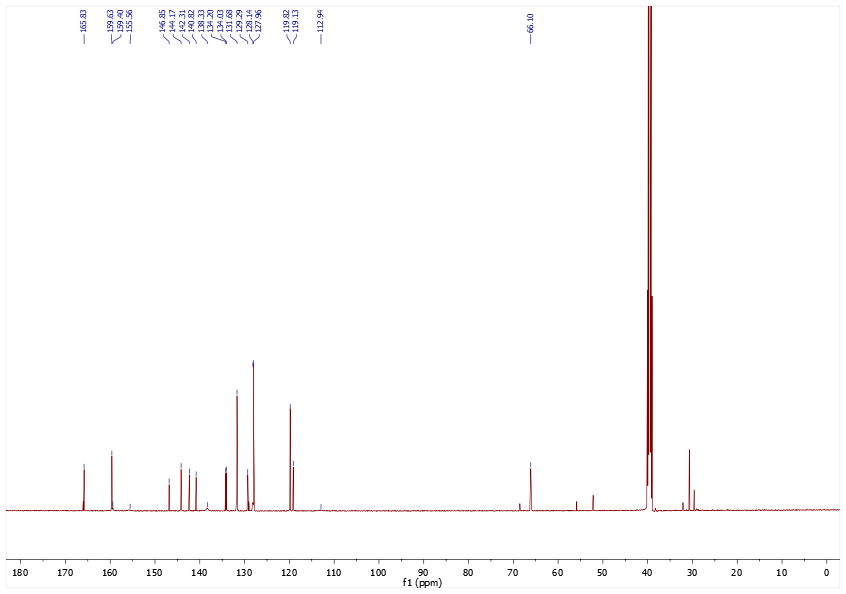

^19^F NMR Spectrum

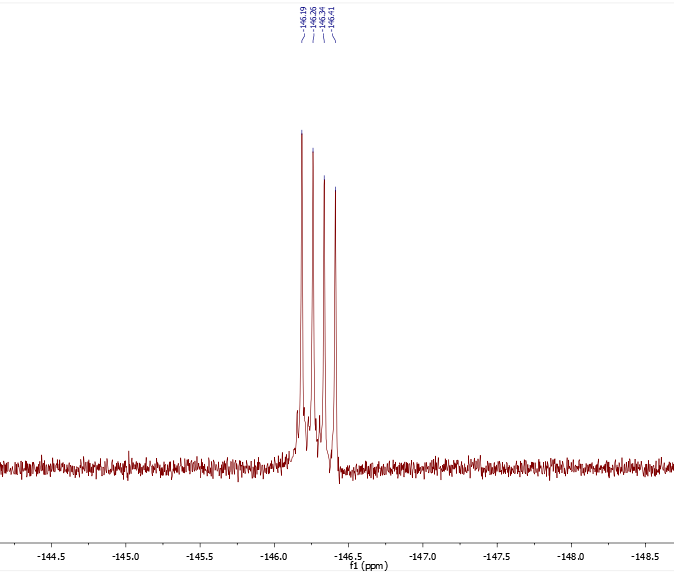
